# Spatiotemporal modeling reveals high-resolution invasion states in glioblastoma

**DOI:** 10.1101/2023.12.05.570149

**Authors:** Varsha Thoppey Manoharan, Aly Abdelkareem, Samuel Brown, Aaron Gillmor, Courtney Hall, Heewon Seo, Kiran Narta, Sean Grewal, Ngoc Ha Dang, Bo Young Ahn, Kata Otz, Xueqing Lun, Laura Mah, Franz Zemp, Douglas Mahoney, Donna L. Senger, Jennifer A. Chan, A. Sorana Morrissy

**Author notes:** Senior co-corresponding authorsl.

## Abstract

Diffuse invasion of glioblastoma cells through normal brain tissue is a key contributor to tumor aggressiveness, resistance to conventional therapies, and dismal prognosis in patients. A deeper understanding of how components of the tumor microenvironment (TME) contribute to overall tumor organization and to programs of invasion may reveal opportunities for improved therapeutic strategies. Towards this goal, we applied a novel computational workflow to a spatiotemporally profiled GBM xenograft cohort, leveraging the ability to distinguish human tumor from mouse TME to overcome previous limitations in analysis of diffuse invasion. Our analytic approach, based on unsupervised deconvolution, performs reference-free discovery of cell types and cell activities within the complete GBM ecosystem. We present a comprehensive catalogue of 15 tumor cell programs set within the spatiotemporal context of 90 mouse brain and TME cell types, cell activities, and anatomic structures. Distinct tumor programs related to invasion were aligned with routes of perivascular, white matter, and parenchymal invasion. Furthermore, sub-modules of genes serving as program network hubs were highly prognostic in GBM patients. The compendium of programs presented here provides a basis for rational targeting of tumor and/or TME components. We anticipate that our approach will facilitate an ecosystem-level understanding of immediate and long-term consequences of such perturbations, including identification of compensatory programs that will inform improved combinatorial therapies.

## Introduction

Glioblastoma is the most common malignant brain tumor in adults – incurable despite multi-modal treatment with maximal safe surgical resection, radiation and chemotherapy^1^. Part of the treatment challenge in GBM stems from its highly invasive phenotype, wherein individual tumor cells move through normal tissues diffusely or follow perivascular routes or white matter tracts to spread far beyond the main tumor mass^2^. Although surgery removes the bulk of the tumor, these infiltrating cells are left behind as they are not only difficult to detect but also impossible to remove without unacceptable neurologic consequences. The residual cells can then continue to evolve and adapt to the selective pressures of conventional and rational therapies – a process that is multifaceted, and that involves genetic heterogeneity, phenotypic plasticity, and the ability to engage with and co-opt the tumor microenvironment (TME) into a pro-tumorigenic state^2–7^ – to result in recurrence. Understanding of the biology of the invasive front and delineating the mechanisms by which these cells engage with their surrounding normal cells and environment to promote the malignant phenotype is thus of high clinical importance.

Multiple factors have hindered our ability to understand the invasive front, however, and its relationships with the rest of the spatial glioblastoma ecosystem. First, surgery removes the most solid tumor tissue, but the outermost regions of infiltration are often inaccessible and undersampled due to clinical limitations. Second, surgery and tissue banking typically yield small and unoriented tissue fragments with an unknown spatial relationship relative to each other. Thus, most GBM literature describes heterogeneity within only small regions of the highly cellular or highly vascularized resectable portion of the tumor and are unable to capture overall tumor organization. Third, studies to date have implicated multiple factors in invasion, including signaling pathways^8,9^, components of the extracellular matrix (ECM)^10,11^, and interactions with nearby or distal cells of the tumor^12,13^ and TME^14^, but there have been limited approaches and tools to contextualize the many processes as part of an ecosystem of interdependent processes within the tumor’s overall organization^15^. These challenges have collectively limited our ability to assess global patterns of adaptation to local tissue contexts, for instance invasion along white matter tracts or perivascular routes within the same tumor.

Fortunately, new technologies for spatial profiling, which can delineate the organization of tumor transcriptional cell states in relation to each other, to genetic diversity, and to metabolic and cellular diversity of the TME, have the potential to greatly increase our understanding of tumor ecosystems^16–20^. Nevertheless, the spatial profiling platforms available to date that offer global transcriptome coverage –and that could therefore support exploratory studies– do not have single cell resolution (NanoString GeoMx, 10X Genomics Visium). The resulting data are therefore mixtures of cell types and states that require computational deconvolution. Strategies to deconvolute spatial data employ either supervised approaches (requiring matched single cell data to infer cellular composition within each profiled region)^21–23^, unsupervised approaches based on matrix factorization or probabilistic modelling^24–27^ (to identify latent gene expression programs representing cell types or states), or semi-supervised approaches^28,29^. Regardless of strategy, most current applications of deconvolution have not robustly disambiguated low frequency signals and have primarily focused on regions where signals of interest are >20% of the total. This excludes areas of diffuse tumor infiltration^16,17,20^, as well as lower frequency cells and states of the TME. Altogether, these limitations hold back key aspects of tumor biology from comprehensive study, including components of the TME and cellular interactions that drive tumor growth and invasion.

Here, we aim to address these challenges by coupling transcriptome-wide spatial profiling of xenografted GBM cells with temporal sampling throughout tumor progression, and an analytic workflow that enables highly specific and sensitive detection of cellular programs (**Fig. 1a**). A key strength of the xenograft strategy is genomic distinction between human tumor cells and mouse-derived TME, which we leverage to boost signal detection in regions with low tumor content. This approach captures transcriptional phenotypes across the whole tumor including areas of invasion. The GBM lines selected represent multiple common genetic drivers, allowing identification of both universal and genotype-enriched transcriptional programs that can be linked back to external patient cohorts. To identify molecular programs, we developed a computational workflow centered on unsupervised deconvolution that does not require matched single cell data and enables *de novo* discovery of both known and novel expression programs corresponding to cell types, cell activities, or combinations thereof. This led to deconvolution of the mouse brain and TME with far greater granularity than previously achieved for spatial data, distinguishing 71 cell types and anatomic structures, and 19 TME-related programs in this cohort. Many of the TME programs (including astrocytic, myeloid, and vascular cell types and states) showed spatiotemporal variation in abundance, reflecting dynamic changes during tumor growth and invasion. We further identified multiple highly resolved human tumor cell programs and explored within-tumor and tumor-TME crosstalk across regions of variable tumor density, identifying unique interactions along distinct routes of invasion. Altogether, our work provides insight into previously elusive aspects of the GBM ecosystem and enabled systematic exploration of the invasive front.

**Figure 1.**
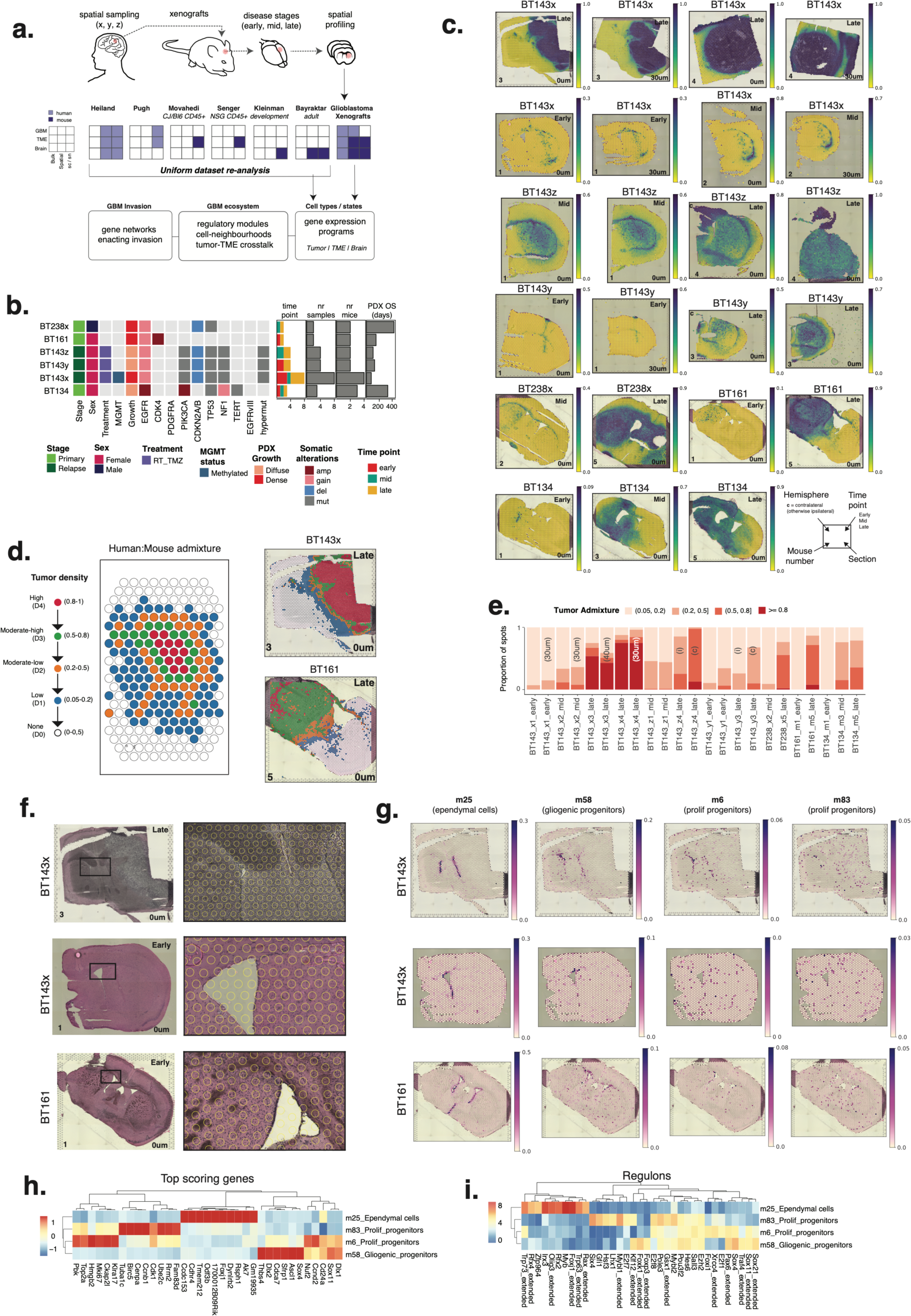
Study overview. **(a)** Schematic representation of data generation and analysis. Cell lines established from patient tumors were xenografted into mice. In some patients, multiple lines were generated from tumor core (x), contrast-enhancing region (y), and leading edge (z). Early, mid, and late timepoints were profiled with the Visium platform, quantifying gene expression programs from tumor, TME, and normal brain using unsupervised deconvolution, their relationship within the GBM ecosystem, and a focus on invasion. Additional available datasets were similarly deconvoluted, to enable comparison between profiling platforms and sample types. **(b)** Genomic overview of the xenograft cohort, including somatic mutations and copy number events in each line, corresponding sample and patient characteristics, and xenograft samples collected. **(c)** Spatial plots of 23 coronal sections showing the ratio of human-mouse transcriptome data. High values (indigo) indicate high tumor density, low values (yellow) indicate low or absent tumor density. **(d)** Illustration of tumor density regions (left panel) stratified based on human:mouse transcriptome admixture, and two selected samples (right panels). **(e)** Proportion of spots in D1-D4 tumor-density ranges per sample (replicate annotations overlaid on plot). **(f)** H&E histology images with overlay of spot perimeters. **(g)** Spatial plots of mouse program usage. Dynamic scales indicate proportional program usage for each spot. **(h-i)** Hierarchically clustered heatmaps of top 10 genes (**h**; rank ordered cNMF-derived gene scores) and top 10 transcription factors (**i**; rank ordered SCENIC activity scores) in mouse programs from panel h.

### stRNAseq profiling and program discovery in GBM xenograft models

To comprehensively capture spatial transcriptional heterogeneity *in vivo*, we selected 6 well characterized brain tumor initiating cell (BTIC) lines^30^. These derive from four patients spanning a range of clinical variables including sex, exposure to standard therapy, MGMT methylation status, and common GBM molecular drivers (**Fig. 1b, Supplementary Table 1a**). In addition to diversity of genetic drivers, our cohort further included BTIC lines from two patients (BT143 and BT238) that captured acquired phenotypic diversity during tumor evolution. In these patients, multiple lines were concurrently derived from anatomically distinct regions of the tumor as defined by pre-operative MRI, including the densely growing core (x), the contrast-enhancing highly vascularized tumor margin (y), and the highly diffuse leading edge of the tumor (z) that is typically outside of the surgical margin. Lines from each patient shared genomic drivers but maintained distinct growth phenotypes *in vivo*, with x lines growing densely relative to the diffusely infiltrating y and z lines (**Fig. 1c**). The consistency of x/y/z growth patterns across patients indicated that predictable phenotypic adaptations can be acquired by GBM cells during tumor evolution, that these adaptations relate to spatial context (e.g., core vs edge), and that they are heritable, likely being fixed in the genome or epigenome. Similar observations have been made for other lines derived from edge versus core^12^ and in the context of adaptation to hypoxia^19^. Our x/y/z BTIC lines thus offer a rare opportunity to investigate expression programs underlying dense versus diffuse growth. Finally, to capture invasion dynamics across tumor stages, our cohort includes early, mid, and late timepoints of growth based on known time to endpoint (ranging between 76-428 days across lines) **(Fig. 1b, Supplementary Table 1a).**

We profiled 51,952 individual spatial transcriptomic RNAseq (stRNAseq) measurements (spots) using Visium, from a total of 23 samples (**Fig. 1c, “Methods”**) spanning all lines and timepoints. In many cases, we were able to fit a complete coronal brain section diagonally within the capture area, ensuring profiling across the whole tumor and invasive front. In other cases, the injection side (i) and contralateral side (c) were mounted on separate capture areas (e.g., BT143y/z endpoint samples, **Fig. 1c**). Further, we also profiled sequential sections cut 30-40um apart but did not observe notable biological variability (data not shown). We used human-to-mouse transcriptome admixture to calculate the relative contribution of tumor cells to the transcriptional output of each spot. Admixture ranged from 0 in spots with 100% mouse cells, to 1 in spots with 100% tumor cells (**Fig. 1c-e, Supplementary Fig. 1a,b**). The sensitivity with which we could distinguish mouse from human cells enabled us to focus our next analyses on regions of high (80-100%; D4), moderate-high (50-80%; D3), moderate-low (20-50%; D2), and low (5-20%; D1) tumor cell density, as well as spots of mouse brain without tumor (D0). We observed highly variable levels of tumor density among lines and timepoints across tumor regions defined in this way. BT143x stood out as the most densely growing line with many spots at >80% tumor density by endpoint (**Fig. 1e**). BT161 and BT238 were the next-most densely growing, with many spots having 50-80% tumor cell density at endpoint. In contrast, BT134 endpoint tumors, and all earlier timepoints across lines grew diffusely.

### Gene expression program discovery in human and mouse cells

Since stRNAseq does not have single cell resolution, we utilized unsupervised deconvolution to identify admixed cell types and states separately for human and mouse data. We used consensus Non-negative Matrix Factorization (cNMF)^31^ to identify robust transcriptional programs across all 23 samples together and quantify their relative usage within spots. Factorization yielded 15 human tumor cell programs (referred to as *h1*-*h15*) and 90 mouse brain and TME programs (referred to as *m1*-*m90*), in line with the greater transcriptional diversity of the mouse brain. To decipher if programs represented cell types, cell states, or anatomical structures, we used gene set enrichment analysis (GSEA) and established reference marker genes for normal mouse brain cell types^28,32,33^, TME-specific cell types^7,28,34^, and human GBM states^4,5,17,35–40^ (**Supplementary Table 2a, Methods**).

Human tumor programs were most diverse in the core, with 3-4 discernable programs per spot in high density regions, and 1-2 programs present per spot in lower density regions (**Supplementary Fig. 1c**). Mouse programs included 11 cell activities and 18 cell types resident in the normal brain, 19 TME-specific or enriched cell types and states, and 42 programs representing combinations of cell types that could not be further disambiguated at this level of factorization (**Supplementary Table 2b**). Annotation by a neuropathologist confirmed that most of these combination programs corresponded to anatomic regions or structures of the mouse brain (**Supplementary Table 2b**). We detected an average of 5 normal brain mouse programs per spot in regions without GBM (D0) and in the tumor leading edge (D1), with diversity decreasing at higher tumor densities (D2-D4; **Supplementary Fig. 1c**). In contrast, only 1-2 TME-related programs were observed per spot in tumor regions. Overall, across both human and mouse data our approach was able to quantify usage of up to 8 programs per spot, showing far greater sensitivity than previously achieved with stRNAseq analyses (i.e., ∼2 programs/spot)^16^.

To ensure our workflow generated meaningful results, we evaluated a set of mouse programs expected to closely match previous literature for cell types and anatomic structures. A first example is program *m25*, which corresponds to ependymal cells based on enrichment of marker genes (**Supplementary Table 2d, 5b**). Ependymal cells form a single-cell layer restricted to the lining the ventricles, and indeed, m25 was localized to the ventricular lining with no background signal elsewhere (**Fig. 1f-g**). Usage values ranged between 0.3-0.5, indicating ependymal cells make up one third to one half of the signal in these spots, in line with known cell composition in the ventricular lining^41^. Analysis of marker genes and transcription factor (TF) activity further validated *m25* as an ependymal program (**Fig. 1i,j**). Within the subventricular zone, we also identified progenitor cell programs at lower cellular frequency per spot (3-10%), including gliogenic progenitors concentrated in the dorsolateral ventricular region (*m58*), and two proliferating progenitor programs (*m6*, *m83*) located in the lateral ventricular lining and with diffuse spread into the brain parenchyma (**Fig. 1f,g**). As a second example, we could distinguish pericytes from endothelial cells (**Supplementary Fig. 1e**), even though these cell types always co-occur spatially within the vasculature, represent a minority of signals within a given Visium spot (max of 8% usage for the pericyte program), and have not previously been deconvoluted in spatial datasets^16,17,20,28^. In a final example, we highlight identification of cortical layers with high resolution (**Supplementary Fig. 1f**), including the outermost meninges (*m4*) and vascular leptomeningeal smooth muscle cells (*m11*). The cortical layers are arranged in a spatially overlapping manner that reflects the changing composition of cell types and states along the radial axis of the brain. This compositional gradient is quantitatively captured by the program usage values, recapitulating the level of spatial overlap between cortical programs (**Supplementary Fig. 1f**). Based on these observations, we conclude that our deconvolution and annotation approach is highly specific (enabling identification of unique cell types, states, and anatomic structures), is highly sensitive (capable of deconvoluting signals with low overall signals in the range of 3-10%), can identify both spatially coherent and diffuse programs, and provides interpretable usage values that quantify up to 8 programs within each Visium spot.

### Tumor programs span progenitor states to invasion phenotypes

Having established the sensitivity and specificity or our deconvolution approach, we characterized 15 *de novo* human tumor cell programs using a similar strategy (**Fig. 2a-d, Supplementary Fig. 2a,b and Supplementary Table 2b**). This included pathway-based and marker gene-based annotations, transcription factor (TF) activity-based similarity, and spatial co-occurrence. These analyses were used to label and assign each program to broader themes based on all information (**Supplementary Data 3a-e**). The 15 human tumor programs are fully described in the context of known GBM transcriptional classes, pathways, and TME spatial neighborhoods in **Supplemental Results**.

**Figure 2.**
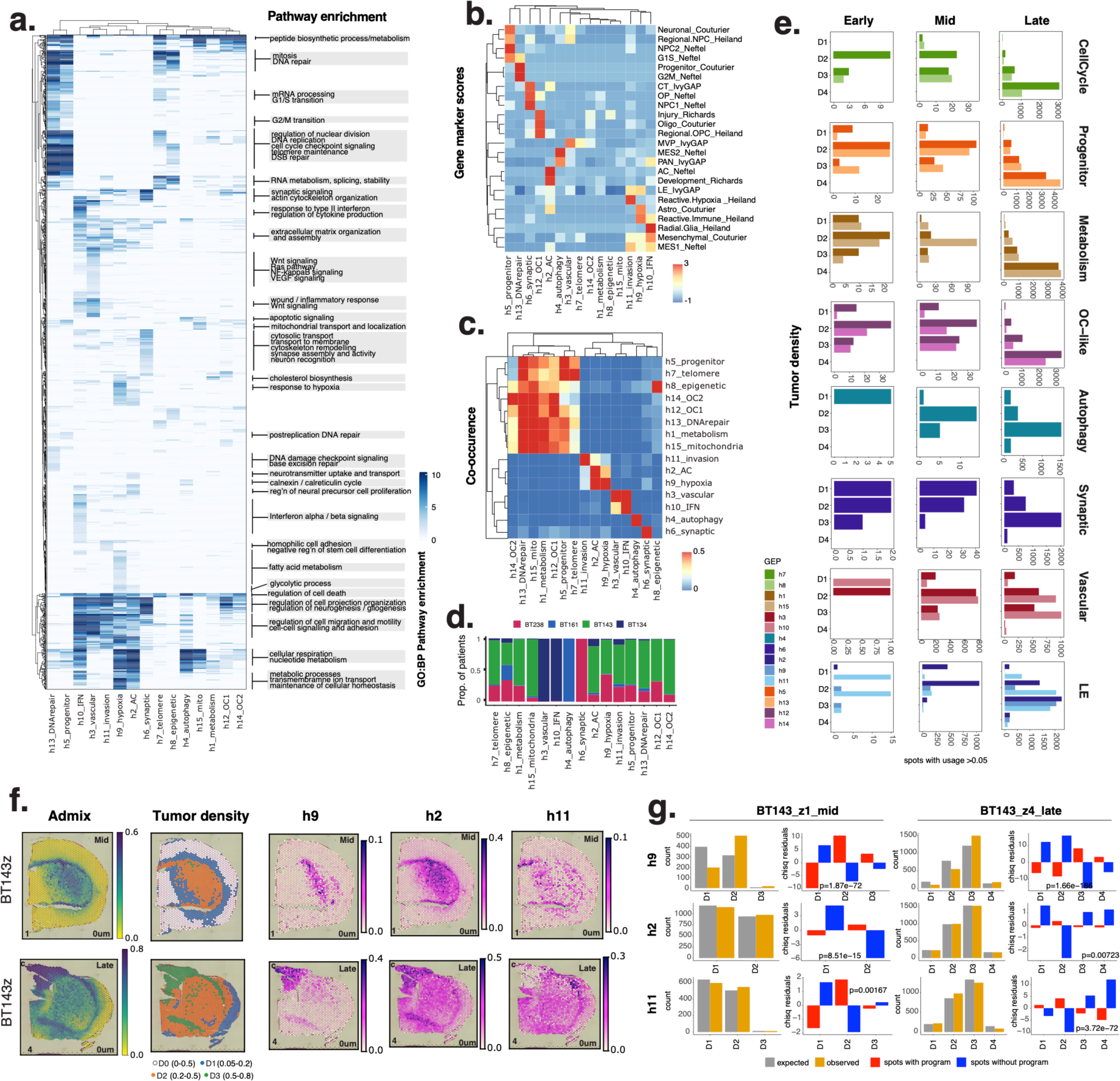
Tumor programs. **(a)** Summary of biological processes (GO:BP; rows) significantly enriched among the top 2000 genes of tumor gene expression programs (columns). **(b)** Heatmap of marker gene enrichment scores of tumor programs (columns) calculated for GBM cell-states (rows) in external datasets. **(c)** Proportion of spatial overlap between TME programs. Spatial overlap for a pair of programs is calculated as the proportion of spots in program 1 that also have usage of program 2, and vice versa. Minimum usage threshold=0.1. **(d)** Proportion of tumor spots per patient with usage (> 0.1) the 15 tumor programs, ordered by category. **(e)** Bar plots of number of spots with usage of each tumor program, stratified by tumor cell density and timepoint. Programs are grouped by thematic category. **(f)** Spatial plots of selected tumor program usage, with tumor admixture and tumor density groups on the left. **(g)** Number of expected and observed spots with usage of selected programs, and chi-squared residuals and pvalues indicating the significance of difference between observed and expected numbers across categories (D1-D4).

We classified programs into 6 major themes, representing progenitor states, cell cycle, metabolism, astrocytic-like, oligodendrocytic-like, and invasion. Progenitor programs (*h5_progenitor*, *h13_DNArepair*) were most prevalent in high tumor cell density regions (D4) (**Fig. 2e, Supplementary Fig. 2c, Supplementary Table 3f**), along with one of the cell cycle programs (*h7_telomere),* indicating that cycling progenitors preferentially reside there. The OC-like programs (*h12_OC1, h14_OC2*) also showed enrichment in denser tumor regions (**Fig. 3e**), with *h14_OC2* representing a genetic subclone within the BT143x cell line (**Supplementary Fig. 2d-g, Supplemental Materials**). Outside of the dense tumor core, a trio of programs formed a gradient of outward tumor expansion reflecting a phenotypic arrangement centering on regions of hypoxia (*h9_hypoxia* most centrally located), extending to regions of more diffuse tumor expansion without hypoxia (*h2_AC*), and finally to invasion into the normal brain (*h11_invason*) (**Fig. 2f,g**). This was reminiscent of hypoxia-centered cellular organization within human tumors^16,42^, indicating our xenograft cohort captures this facet of tumor biology, and importantly, extends the previous work by revealing a program of diffuse invasion (*h11*). The *h11_invasion* program scored highly for leading edge gene-sets (LE_IvyGAP; **Fig. 2b**), comprised a majority of D1-D2 spots at early to late timepoints, and was the only tumor program with significant over-representation in D1 regions within individual patients (**Fig. 2g, Supplementary Table 3g**). Finally, we noted that a few programs did not have cohort-wide usage (**Fig. 2d**). Instead, these were prevalent within individual patients suggesting that specific tumor genotypes could be linked to unique repertoires of tumor cell programs, and that larger xenograft cohorts will likely reveal additional insights. In support of this, previous work establishing that EGFR plays a role in errant neovascularization^42^ was in line with enrichment of angiogenic tumor cell programs (*h3*, *h10*) in the tumor line BT134, which harbours high-level EGFR amplification (93 copies).

**Figure 3.**
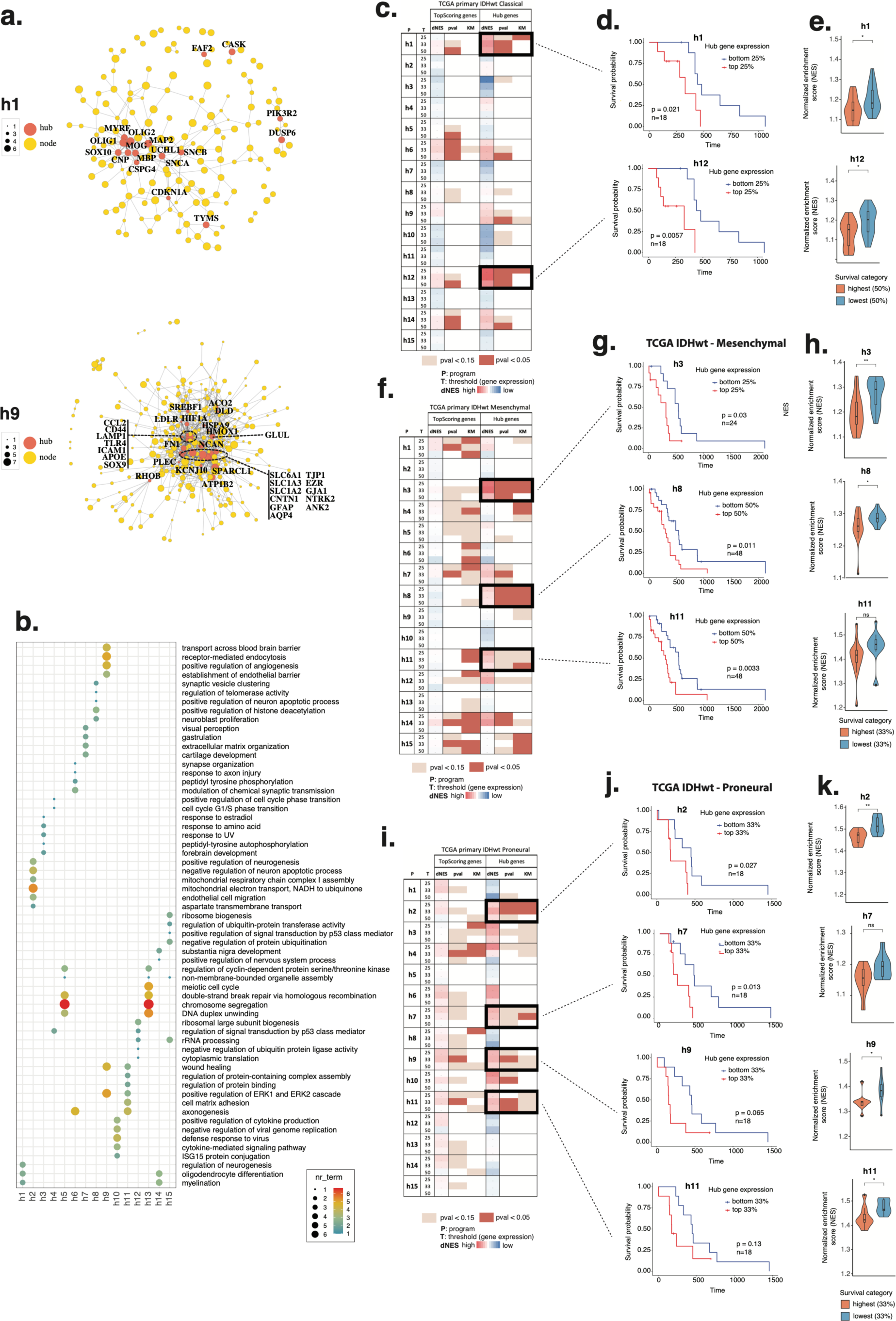
Survival associations. **(a)** Network plot of top-scoring genes (yellow nodes) for selected programs, with hub genes in orange, and labelled with gene name. Node size is proportional to the correlation between a gene’s usage in a program to its expression, and edges represent the functional associations. **(b)** GO:BP terms enriched in hub-genes of tumor programs. **(c,f,i)** Survival results table for Classical IDHwt tumors. For each program, top-scoring program genes and hub genes were tested separately using two methods. First, ssGSEA was run for each geneset, and patients were ranked by survival. Three thresholds were used to separate top and bottom scoring patients (25%, 33%, and 50%). The NES difference between patients with low and high survival (dNES) and the t-test pvalue are reported in separate columns. Second, Kaplan Meyer analysis was performed and pvalues are also reported in a column, colour-coded to highlight blocks of significant and marginally significant pvalues across thresholds and the two testing strategies. **(d,g,j)** Kaplan Meyer plots for selected program in each GBM tumor subtype. **(e,h,k)** Distributions of NES values in patients ranked by survival are shown along with adjusted Student’s t-test significance. (*: p <=0.05; **: p <= 0.01).

Finally, we gauged how prevalent xenograft tumor programs were in other datasets by comparing them with similarly identified programs in external cohorts of human tumors^5,17,30^, brain tumor initiating cell lines^30^, and xenografts^30^ (**Supplementary Fig. 2i, Supplementary Table 3h**). This comparison showed highest correlations for progenitor, cell cycle, and metabolism programs among datasets. We saw less concordance for genotype-specific programs outside of the TFRI cohort (which included the bulk RNA-seq of these xenograft samples) indicating that these programs are less prevalent across GBM patients.

### Tumor program association with survival and transcriptional subtypes

We next asked how the 15 human programs related to survival differences, anticipating that associations may be context-dependent with respect to the GBM transcriptional subtypes (classical, mesenchymal, proneural). For each tumor program we first identified the genes most strongly contributing to program identity (i.e. top-scoring genes) (**Fig. 3a, Supplementary Fig. 3, Supplementary Table 4a**), then performed network analysis to select the subset acting as network hubs based on protein-protein-interactions (**Fig. 3a, Supplementary Fig. 3, Supplementary Table 4b**). Hub genes were distinct among programs (**Fig. 3b**) and well aligned with previously described program themes (**Fig. 2a**). We performed survival analyses in the TCGA cohort using both top-scoring and hub genes and found that hub genes had a strong associations with survival across all transcriptional subtypes (**Fig. 3c,f,i, Supplementary Table 4e**). This was true in both a classic survival analysis (comparing survival of patients ranked by gene-set expression; **Fig. 3d,g,j**), and when comparing gene-set enrichment in patients first stratified by survival outcome (**Fig. 3e,h,k**). Several programs emerged as robustly associated with survival in a subtype-specific manner. For instance, high expression of *h1_metabolism* and *h12_OC1* was robustly associated with poor survival in classical tumors. The GSEA normalized enrichment scores (NES) of *h1* and *h12* genesets were very highly correlated across the TCGA cohort (cor=0.85; **Supplementary Table 4f,g**), indicating that OC-like cells enacting *h1* metabolic activities (primarily cholesterol biosynthesis) operate as a unit, together influencing cell phenotypes related to survival. Mesenchymal tumors were stratified by cell cycle/epigenetic (*h8_epigenetic*) and invasion (*h11_invasion*) programs. Lower NES correlations between *h8* and *h11* (cor=0.3; **Supplementary Table 4f,g**) indicated these programs independently influenced survival. Proneural tumors were also stratified by cell cycle (*h7_telomere*), and the three programs encompassing the hypoxia-to-invasion gradient (*h9_hypoxic, h2_AC1, h11_invasion*). NES correlation values among these programs in TCGA followed the same graded pattern of co-occurrence observed in xenografts (*h9*-*h2*-*h11*), strongly supporting their spatial and phenotypic relationship in human patients (**Supplementary Table 4f,g**).

Having established prognostic value of the tumor programs, we more deeply investigated how these related to known GBM molecular transcriptional states (Neftel)^4^. This was specifically motivated by the weaker observed match of the strongly prognostic invasion programs (*h2*, *h11*) to NPC-like states (**Fig. 2b**), despite previous links between NPC-like states and GBM invasion^4,6,43^. Given that multiple states typically co-exist within a tumor and that state transitions can reflect adaptations to the TME^4,6,37^, we anticipated correspondence between Neftel states and tumor density. As expected, each xenograft harbored all four states (**Supplementary Fig. 4a-c**), with NPC-like and MES states less abundant overall, but with significantly higher prevalence in D1-D2 regions relative to the tumor core (**Supplementary Fig. 4a**). We also found that tumor-baseline (majority) states were evident in each xenograft sample, were stable across timepoints and tumor density regions, were independent of cNMF program usage, and therefore likely intrinsic to each line, as previously observed^6^ (**Supplementary Fig. 4c**). By then calculating the rate of transition away from tumor-baseline in spots with usage of individual programs, we established that transitions were common across programs (**Supplementary Fig. 4d**). This supported the high plasticity of Neftel states among the tumor programs defined here, and specifically for *h11_invasion* in regions of diffuse infiltration (D1) where dynamic transitions from an AC baseline to an NPC-like state predominated across all timepoints. This supported *h11* as an invasion program largely orthogonal to the previously established NPC-like state.

### Microenvironment programs encompass cell types and states

Of the 90 mouse programs, we focused on 19 that were either specific to or highly enriched in regions of tumor, each with dynamic spatial and temporal kinetics. These TME-programs were broadly categorized into 11 cell types and 8 cell activities based on marker gene, TF-activity, and pathway annotations, and co-localization (**Fig. 4a-d, Supplementary Fig. 5a-c, Supplementary Table 2b,5a-d, 3e**). Programs were labeled as cell types when marker gene-based annotations were unambiguous and strong, otherwise, labels thematically reflected highly enriched pathways indicating cell activities. All programs of the TME were detected in all lines (**Fig. 4e**). We describe the resulting *in vivo* spatial xenograft TME catalogue in **Supplemental Results**, covering astrocytes, vasculature, immune cells (microglia, monocytes, macrophages), and multiple activity programs.

**Figure 4.**
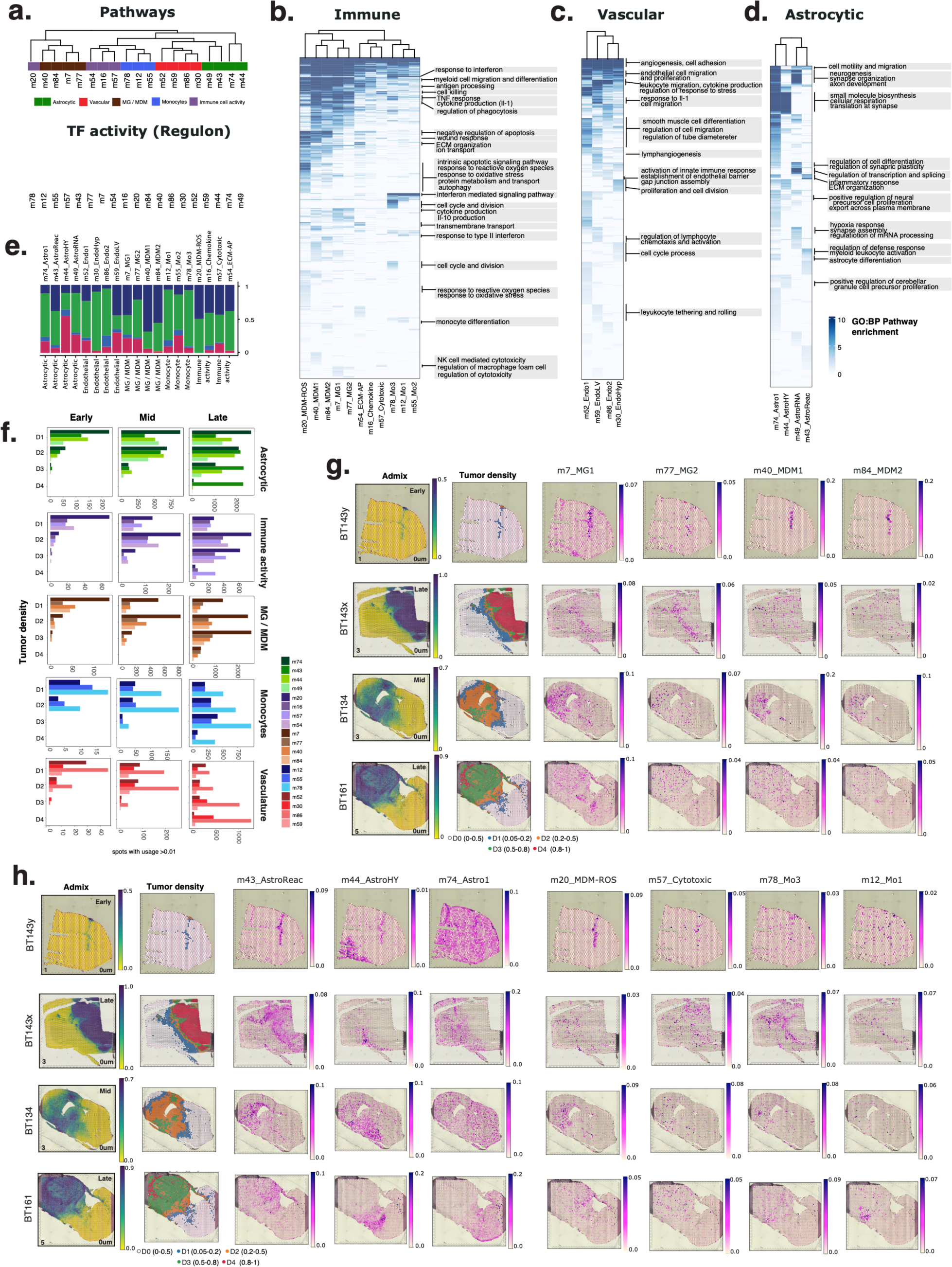
TME programs. **(a)** Pathway-based hierarchical relationship of 19 TME programs based on clustering of significant GO:BP gene-sets (adjusted p.value < 0.05) enriched among the top 2000 genes in each program (ranked based on program gene scores) (top panel). Hierarchincal clustering based association of programs based on TF-activity scores (bottom panel). **(b-d)** Pvalue heatmaps and annotations of **(b)** immune, **(c)** vascular and **(d)** astrocytic programs. **(e)** Proportion of tumor spots per patient with usage (> 0.05) the 19 TME programs, ordered by category. **(f)** Bar plots showing the number of spots with usage of each TME program, stratified by tumor cell density and timepoint. **(g-i)** Spatial plots of **(g)** immune, **(h)** astrocytic and **(i)** activity programs in selected samples, with tumor admixture and tumor density indicated on the left for reference.

TME programs were broadly organized along a spatial axis corresponding to tumor density. For instance, we observed that mature astrocytes (*m74_Astro1*) widespread and resident throughout the normal mouse brain were prevalent in the invasive tumor front but excluded from dense tumor regions (**Fig. 4f,h**). *m49_AstroRNA*, representing normal astrocytic activities (related to RNA processing, proliferation, glutamatergic synapses) was prevalent in astrocytes localized to the normal brain and tumor periphery (**Supplementary.** **Fig 5d, Supplementary Table 5e**). Within the context of the tumor however, a dramatic state shift toward a reactive program (*m43_AstroReac*) was enacted across the full spectrum of tumor density and sustained at all timepoints of disease progression (**Fig. 4f, Supplementary Fig. 5d, Supplementary Table 5e**). Astrocytes in this reactive state were enriched for terms relating to proliferation, inflammatory response, and ECM organization, and had a strong match to established reactive signatures^44^ (**Supplementary Fig. 5e**). Altogether, these state-change patterns supported phenotypic co-option of normal astrocytes along the advancing tumor front. We were surprised to also observe tumor-association of a regional astrocytic cell subtype or activity (*m44_AstroHY*), characterized by pathway enrichment of terms relating to precursor cell proliferation. This seemingly homeostatic program, highly used within hypothalamic regions at all timepoints (**Fig. 4h**), also showed early and sustained tumor-enrichment (**Supplementary Fig. 5d, Supplementary Table 5e**), indicating that tumor-proximal astrocyte transitions to this state could play an important a role within the context of the GBM TME.

**Figure 5.**
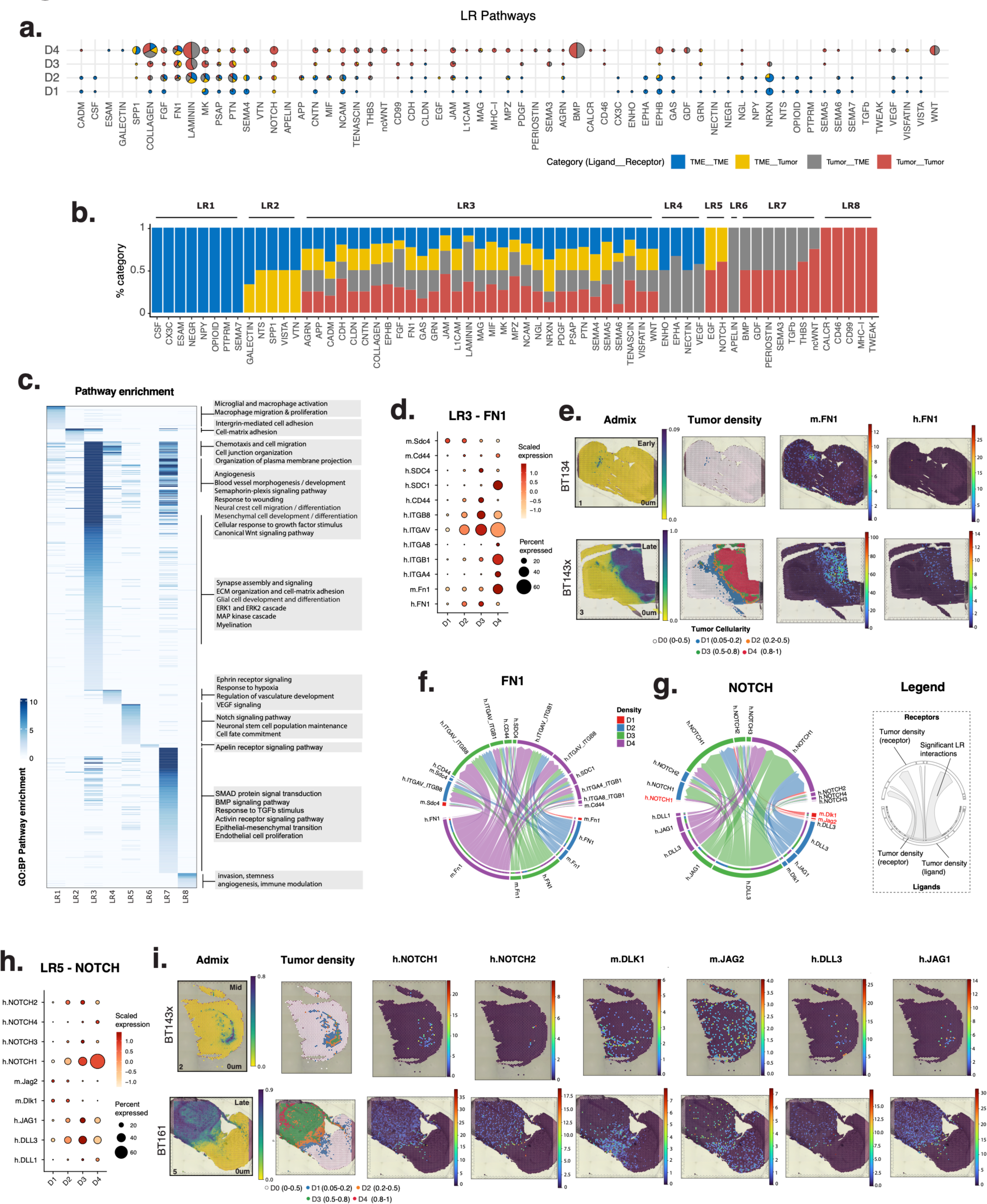
LR communication. **(a)** Significant pathways (x axis) from CellChat are shown for each tumor density group (y axis). Piecharts indicate the proportion of human and mouse ligand-receptor interaction types. Piechart size represents the total number of interactions active in a pathway. **(b)** Stacked barplot indicating the proportion of interactions per pathway identified as significant. LR interactions are categorized and colored based on the species of the ligand and receptor involved, resulting in 4 interaction categories (ligand receptor): TME TME, TME Tumor, Tumor TME, and Tumor Tumor. Pathways are categorized into 8 groups (LR1-LR8) based on combinations of the 4 interaction categories. **(c)** Overview of significantly enriched biological processes among ligand and receptor genes in each LR group. **(d,h)** Scaled expression levels (z-scores) of ligand and receptor genes per pathway, stratified by tumor density (D1-D4). **(e,i)** Spatial plots of human (h.GENE) and mouse (m.Gene) genes in selected samples (right panels) with tumor admixture and tumor density for reference (left panels). **(f,g)** Chord diagrams show receptor (bottom) and ligand (top) interactions for the FN (g) and NOTCH (h) pathways. Interactions are analyzed between all possible pairs of tumor-density groups which make up the sources (outer, lower semi-circle) and targets (upper semi-circle), with links representing the signaling strength of interactions between them (i.e. communication probability). Text labels of D1 interactions between human NOTCH1 and mouse ligands are in red.

We observed similar regional specificity of endothelial programs, with normal vasculature in the normal brain and invasive front (*m52_Endo1*, *m59_EndoLV*) gradually replaced by tumor-enriched vascular programs (*m86_Endo2*, *m30_EndoHyp*) within the denser and more hypoxic regions of tumor (**Fig 4c,f**). Microglial programs were also present diffusely throughout the normal brain (**Fig. 4f,g**), had an early response to sites of injury (injection tract) and persisted long-term within lower density tumor areas (D2-D3) (**Fig. 4f, Supplementary Fig. 5d, Supplementary Table 5e**). Of these, *m7_MG1* was more abundant overall, scoring strongly for response to injury and antigen processing and presentation, while *m77_MG2* was involved in apoptotic cell clearance (**Supplementary Table 5a**), indicating these microglial subpopulations play different roles within the tumor. Two monocyte derived macrophage (MDM) programs were rapidly recruited to early lesions and were otherwise absent from the normal brain (**Fig. 4f,g**). Although these two MDM programs had broadly similar spatial distributions, they enacted distinct activities possibly related to their micro-local spatial contexts – *m40_MDM1* was enriched in terms relating to cell-matrix adhesion, chemotaxis and migration, while *m84_MDM2* scored highly for macrophage proliferation and phagocytosis (**Fig. 4b, Supplementary Table 5a**). Of the multiple immune cell activities identified (**Supplemental Results**), we highlight the cytotoxic program *m57_Cytotoxic* as primarily enacted by MG and MDMs (based on co-localization; **Supplementary** Fig. 5c**, Supplementary Table 5d**), and observed as the most prevalent immune activity in D4, indicating that cytotoxicity plays a more central role in denser regions (**Fig. 4f,i**).

We sought to compare this highly detailed catalogue of TME programs with programs identifiable in external cohorts. We used cNMF to similarly identify programs in 5 additional datasets, including single cell and spatial data from non-tumor-bearing developing and adult mouse brain (Kleinman, Bayraktar)^28,32^, and single cell data from CD45+ cells sorted from syngeneic (Movahedi)^34^ and xenograft (Senger)^7^ GBM models (**Supplementary Fig. 5f, Supplementary Table 5f**). After factorizing these datasets, we identified the best match between the TME programs and the resulting ensemble of cNMF solutions, observing general agreement along expected themes. For instance, astrocytic and endothelial programs had higher correlation with programs in normal brain datasets compared to CD45+ enriched datasets where non-immune cells were depleted. Interestingly, we saw better matches between *m52_Endo1* and Visium rather than single-nuclei data (Bayraktar), indicating possible loss of normal vascular cells during single cell sample processing. Brain-resident microglial programs were also more prevalent in normal brain versus GBM datasets, whereas the opposite was true for MDM programs. Monocytes also showed specific enrichment in the GBM datasets, as expected from their recruitment during tumor progression. Overall, broad concordance among cohorts and data types indicated that the TME program catalogue defined here represents meaningful biological signals.

### Tumor-Microenvironment crosstalk

Cellular communication is vital to the tumor ecosystem, with cell-cell contacts, tumor-extracellular matrix (ECM) interactions, and secreted signaling all pivotal to tumor growth, adaptation, and invasion. To better understand how the glioma and non-malignant cells described here communicate across distinct niches, we quantifed directional ligand-receptor (LR) signaling using CellChat^45^, expecting to capture distinct interactions in dense versus invasive regions reflective of cellular changes in TME composition. We surveyed all possible tumor-tumor, tumor-TME, and TME-TME interactions, stratified by density region (**Fig. 5a, Supplementary Fig. 6a**). In all, 63 pathways were involved in significant crosstalk, comprising 8 major groups based on the directional involvement of human versus TME ligands and receptors (LR1-LR8; **Fig. 5b**). Within the TME (LR1) for instance, we observed that CSF-CSF1R signaling in D1-D2 regions involved microglia as the primary signal-receiving cells, based on high scores for CSF1R in these programs (CSF1R gene rank *m7*=16, *m77*=22; **Supplementary Table 6a**). In LR2, significant Spp1-CD44 crosstalk took place between macrophages and reactive astrocytes at early timepoints, and also between macrophages and tumor cells at later stages (**Supplementary Fig. 6b-e, Supplementary Table 6a**). In this case, the spatial data enabled distinction of mouse versus human receivers based on spatial co-localization of mouse sender and receiver cells at early timepoints (**Supplementary Fig. 6e**). LR8 involved within-tumor signals related to invasion and stemness. In this group, CALCR signaling stood out based on the CALCRL receptor as the top scoring gene in the *h5_progenitor* program (**Supplementary Table 6a**). CALCRL had previously been linked to glioma cell proliferation, has been negatively associated with glioma prognosis, and promotion of angiogenesis^46^. Based on these associations, we speculate that the observed enrichment of the *h5_progenitor* program along with tumor-enriched vasculature programs in D4 could be related to vascular stem cell niche development and maintenance. Indeed, CALCRL is a marker of stemness and a promising candidate therapeutic target in other diseases^47^.

By far the most abundant group of pathways encompassed multidirectional signaling within and between tumor and TME (LR3), including MK and PTN pathways, AGRN, JAM and NCAM (cell-adhesion), tumorigenic NOTCH, PDGF, FGF, and angiogenic VEGF (**Fig. 5a,b, Supplementary Table 6b,c**). We noted that multiple LR3 pathways converged on formation and maintenance of the ECM (tenascin, laminin, collagen, fibronectin). Laminin and tenascin ligands were primarily human (LAMB2, LAMA4, TNC, TNR), while fibronectin ligands were made by both mouse and human cells (**Fig. 5d-f**). Mouse (but not human) Fn1 predominated at early timepoints (**Supplementary Fig. 6d**) and in D1 regions (**Fig. 6f**), providing the majority of this tumor-ECM component in areas of diffuse infiltration. Since vascular programs were strongly associated with Fn1 (*m52, m30, m86*), we conclude that a major invasion route in D1 is along vessels. Collagens also derived from both tumor and TME components, with higher contribution from tumor (**Supplementary Fig. 6a-c, Supplementary Table 6c**). Deposition of human collagens was spatially distinct, with COL6A1 dominant in D1-D3 and specifically associated with the *h11_invasion* program. In D4, COL9A2 and COL9A3 were more prevalent and associated with *h1_metabolism* and *h7_telomere*, indicating changing ECM deposition based on density and niche (**Supplementary Fig. 6b,c**). Mouse collagens Col4a1 and Col4a2 were highest in D4 and derived from the tumor-associated vasculature (with high program scores in *m30_EndoHyp* and *m86_Vasc2*). Altogether, cells of both tumor and TME differentially contributed to the tumor ECM, resulting in a dynamic structural scaffold underlying gradients of invasion and forming the basis for distinct tumor cell niches.

**Figure 6.**
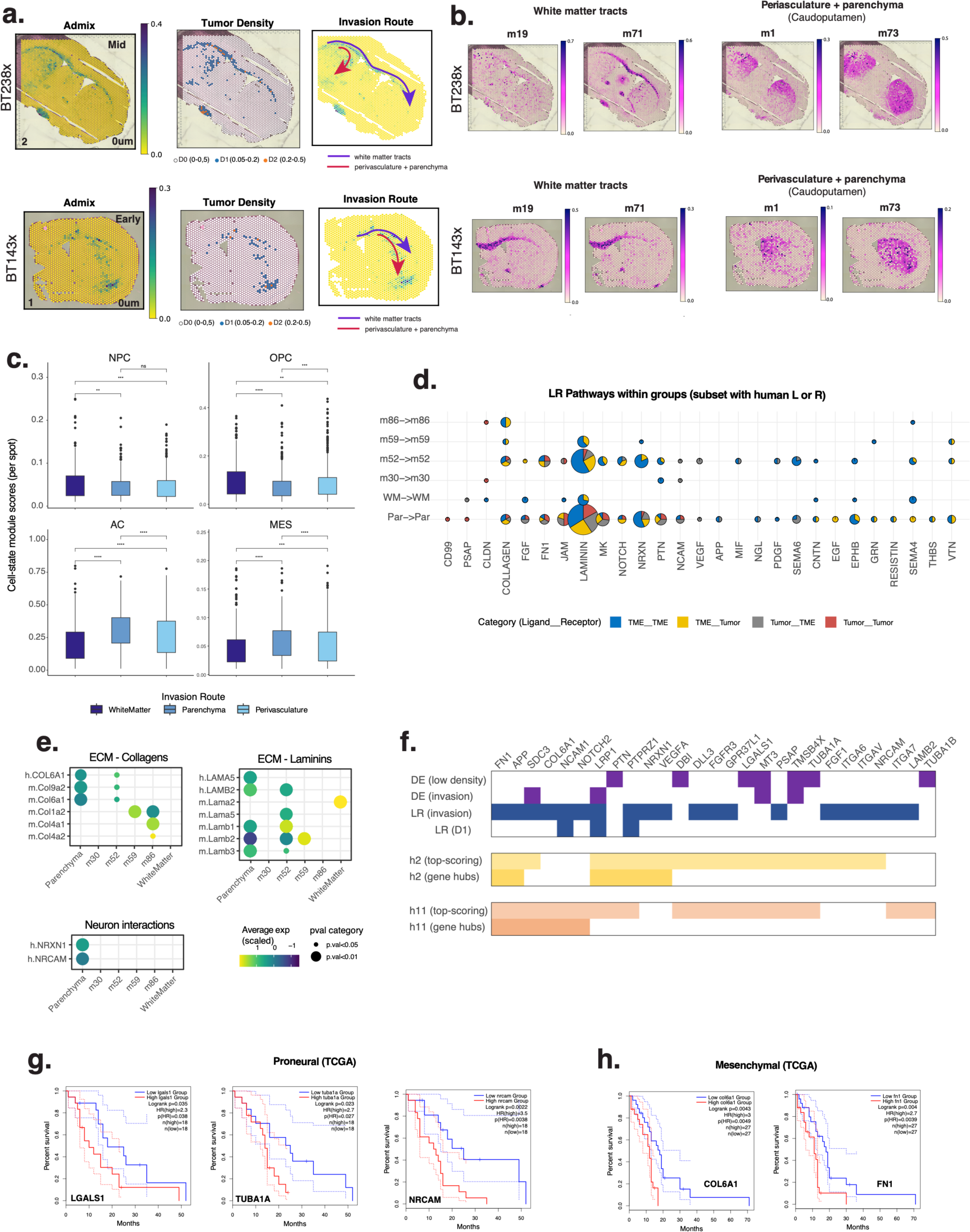
Invasion. **(a,b)** Spatial plots of usage of programs classified either as white matter, or caudoputamen, which includes both vascular and parenchymal invasion routes, with admixture, tumor cellularity, and invasion routes of interest for reference. **(c)** Boxplot of module scores per spot for each GBM cell-state in white matter, parenchymal or perivascular routes of invasion. Significant differences are annotated (wilcoxon rank-sum test ; ns: p > 0.05; *: p <=0.05; **: p <= 0.01) **(d)** Significant pathways (x axis) from CellChat are shown for each D1 category group (y axis), for the subset of pathways with at least one human ligand or receptor. Piecharts indicate the proportion of human and mouse ligand-receptor interaction types. Piechart size represents the total number of interactions active in a pathway. **(e)** Scaled expression levels of selected genes involved in ECM (Collagens and Laminins) and neuronal interactions along white matter, parenchymal or perivascular (m30, m52, m59, m86) invasion routes. **(f)** Summary of invasion-associated genes (columns) based on multiple assessments (rows), including differential expression (DE) between high and low density regions (DE (low density)), DE between parenchyma and white matter spots (DE (invasion)), ligand-receptor interactions from panel (d) (LR (invasion)), and ligand-receptor interactions in D1 from Figure 5a (LR (D1)). **(g,h)** Kaplan Meyer survival curves of genes from panel (f) with prognostic significance in Proneural (e) and Mesenchymal (f) patient tumors.

Within areas of diffuse infiltration (D1) we observed significant invasion-associated midkine signaling through human receptors PTPRZ1 and LRP1, both highly scoring genes in *h11* and *h2* programs. These receptors responded to midkine ligands from astrocytic sources (Mdk highly scoring in *m74_Astro1* and *m44_AstroHY*). Notch signaling also stood out as a main source of mouse-human crosstalk in D1 (LR5; **Fig. 5g-i, Supplementary Fig. 6d, Supplementary Table 6a**). Notch1 is important in GBM invasion along white matter tracts^8^ and for GBM cell survival within the perivascular niche^9^. Our data supports that in D1 regions, the human NOTCH1 receptor (highly scoring in *h2* and *h11*) can bind to mouse ligands Dlk1 and Jag2. These ligands were associated with normal vasculature (Dlk1 in *m52_Vasc1*) and hypothalamic astrocytes (Jag2 in *m44_AstroHYP*), highlighting these cellular programs as distinct TME participants in the invasion signaling axis.

### Molecular and structural contributors to invasion along distinct routes of tumor cell travel

Invasion involves tumor cell movement along white matter tracts, along perivascular routes, as well as directly through the brain parenchyma^2,8,9,48^. We attempted to identify and characterize these routes through association of mouse brain programs with tumor invasion, followed by CellChat^45^ and differential expression analyses^49,50^. We first assessed over-representation of all 90 mouse brain programs in D1 spots, expecting to see enrichment of invasion-relevant cell types or anatomic regions (**Fig. 6a,b**). Indeed, white matter (WM)-related programs had the highest association with D1 tumor regions (**Supplementary Fig. 7a, Supplementary Table 7a**). We therefore selected D1 spots with high usage of mouse WM programs as representative of human tumor cells traveling along white matter tracts. Likewise, we observed that usage of caudoputamen programs (*m1*, *m73*) were also significantly enriched in D1 regions (**Fig. 6a,b, Supplementary Fig. 7a**). Further analysis showed that many caudoputamen spots also had usage of vascular programs (expected from the high diversity of programs identifiable per spot; **Supplementary Fig. 1a**); we therefore further stratified caudoputamen spots based on co-usage of the four vascular programs (**Supplementary Fig. 7b**). This strategy distinguished the D1 tumor cells travelling along each type of vasculature (perivascular routes), from those moving directly through the brain parenchyma (i.e. caudoputamen spots without vasculature). We observed that NPC-like and OPC-like states were more common in D1 tumor cells travelling along white matter tracts, while those within the brain parenchyma were skewed toward MES and AC-like states (**Fig 6c**). Thus, although *h11* and *h2* invasion programs were highly plastic (**Supplementary Fig. 4a-d**), cell state decisions in the invasive front reflect adaptation to the local cellular context of specific routes.

We conducted another CellChat analysis on the resulting 6 groups of D1 spots: white matter (WM), parenchyma (Par), and perivascular routes (*m52*, *m30*, *m86*, *m59*) (**Fig. 6d, Supplementary Fig. 7c-e**). Collagen, fibronectin, and laminin interactions were significant and prevalent along all routes (**Fig. 6d**). Human integrin receptor interactions with various mouse collagens were largely stratified by vasculature programs, suggesting that tumor cells co-opted existing collagen scaffolds within the TME for movement. A notable exception involved human COL6A1 interactions in parenchyma and *m52* regions, indicating that tumor cells specifically required and secreted this ECM component (**Fig 6e**). Similarly, tumor cells bound to multiple mouse laminins (**Fig 6e; Supplementary Table 7b**), but also contributed a select subset of ligands to the ECM (LAMA5 in the parenchyma, and LAMB2 in the vasculature) (**Fig 6e**). A single laminin was specific to the WM route (mouse Lama2) (**Fig 6e**). This gene was highly ranking in program *m71* (newly forming oligodendrocytes), indicating that this oligodendrocyte subtype contributes to the migration scaffold. Collectively, these results highlighted that TME-derived ECM could differentiate between invasion routes, and notably, that a subset of regionally-specific components originated from the tumor cells themselves – indicating the importance of these molecules to glioma cell movement along distinct routes of travel.

CellChat analysis also revealed several signaling interactions specific to the parenchyma, including NRXN, CD99, PSAP, PTN, MK, CNTN, and NOTCH (**Fig. 6d, Supplementary Table 7b**). Of these, we highlight tumor (NRXN1) crosstalk with neuron-derived synaptic adhesion molecule neuroligins (Nlgn1, Nlgn2, Nlgn3) (**Fig. 6d,e**). This previously described mitogenic signaling axis co-opts excitatory neuronal activity toward tumor growth, suggesting that active neurons are playing a role in glioblastoma invasion through the parenchyma^14,51^. In addition, we also further observed interactions between tumor cells via the NRCAM (neuronal cell adhesion molecule) receptor and the mouse Cntn1 ligand – a gene highly scoring in cholinergic and GABAergic neurons, suggesting that these neurons may play additional roles in invasion beyond excitatory activity (**Fig 6e, Supplementary Data 2d**).

As an orthogonal approach to identification of invasion-relevant genes, we performed differential expression analysis between spots with high versus low tumor density in each line (i.e., a program-agnostic approach; **Methods**), and shortlisted genes with recurrent upregulated expression in areas of low tumor density (**Supplementary Table 7c).** Moreover, to further distinguish between invasion routes we performed differential expression analysis between WM and the other 5 groups of interest (parenchyma, *m52*, *m30*, *m59*, *m86*), shortlisting significant genes (**Supplementary Table 7d,e**). We observed high overlap of these differentially expressed genes with the top-scoring and hub genes of programs *h2*, *h9*, *h11* (and to a lesser extent the genotype-specific *h6* and *h10* programs) (**Supplementary Fig. 7f**)

Finally, we combined the evidence from all analyses focused on D1, including the full CellChat analysis (**Fig. 5a**), the invasion route-specific CellChat analysis, and the two differential expression analyses above. Intersecting the resulting genes with tumor programs revealed an overlap of 27 genes that were also top-scoring or network hub genes within the prognostic invasion-related programs *h2* and *h11* (**Fig. 6f**). Nine of these were hub genes in either *h2* (PTN, PTPRZ1, LRP1, NRXN1, VEGFA) or *h11* (SDC3, COL6A1, NCAM1, NOTCH2), while 2 genes were hubs in both (FN1, APP) (**Fig. 6f**). Moreover, in the TCGA cohort, three of these genes were independently associated with poor survival outcomes in Proneural subtype tumors (TUBA1A, NRCAM, LGALS1; **Fig. 6g**). NRCAM is a neural adhesion molecule participating in cell proliferation, axon growth, and synapse formation during neural development, and was previously observed to be overexpressed in brain tumors^52–54^. TUBA1A is a microtubule subunit with effector functions in neuronal migration and causally implicated in neurodevelopmental defects^55,56^. LGALS1 (Galectin-1) has a role in GBM invasion through modulation of cell-cell and cell-matrix interactions^57^ and also promotes an immunosuppressive TME^58^. Another two genes were independently and significantly associated with poor survival in TCGA Mesenchymal tumors (COL6A1, FN1; **Fig. 6h**). Both were previously associated with increased deposition in the ECM by CD133+ glioma cancer stem cells relative to differentiated glioblastoma cells^59^ and COL6A1 has been observed in perivascular and PAN regions in patient samples^60^. Our results now place these genes in relation to each other and to other genes within modular networks that constitute prognostic invasion programs.

## Discussion

In this study we used global transcriptional profiling and unsupervised reference-free deconvolution to perform *de novo* discovery of cell types and states within the GBM ecosystem. We collated a comprehensive and in-depth catalogue of 15 tumor cell programs within the spatiotemporal context of 90 mouse brain cell types, activities, and anatomic structures. The xenograft platform was critical in providing the resolution needed to study the invasive front, where tumor cells were a small minority of the transcriptional output per spot. It has long been recognized that gliomas display histologically distinct patterns of growth relative to normal brain structures. As reported first by Scherer in 1938^48^, these include close interactions of glioma cells with neurons, close interactions with vessels, invasion along white matter tracts, and subpial accumulation. Our findings provide a glimpse into the underlying mechanisms of tumor-microenvironmental cell interactions that support those routes of migration. Importantly, while some of the genes highlighted here had been individually studied in previous work, our deconvolution approach was able to frame these within protein-protein interaction modules that relate expression programs to *in vivo* phenotypes. In particular, genes serving as program network hubs were highly prognostic, stratifying patients across transcriptional subtypes. Whether effective targeting of these prognostic programs could be achieved through perturbation of single or multiple hubs will require future functional testing *in vivo* to ensure that dependencies between invasion programs and invasion routes are faithfully maintained.

In addition to human tumor cell programs, we provide a spatiotemporal breakdown of the glioblastoma TME, encompassing multiple cell types and states. We anticipate that this catalogue will represent a tractable system for gauging the potential impact of rational therapies. For instance, both MDM programs (*m40_MDM1*, *m84_MDM2*) showed high program scores for Lilrb4a – a gene with established roles in promoting immunosuppression, and the target of immunomodulatory therapies currently under development^61^. Similarly, Gpnmb was highly scoring in both MDM programs, has been implicated in proneural to mesenchymal state transitions in GBM^62^ and is another promising TME-specific target in GBM. Our data suggest that targeting of these molecules could effectively impact MDMs but not microglial or monocyte programs, where these genes do not significantly contribute to program identity. More generally, coupling spatial profiling analysis as described here with *in vivo* preclinical assessment of targeted therapies toward either tumor or TME components would facilitate an ecosystem-level understanding of the immediate and long-term consequences of such perturbations. This would include identification of compensatory programs and build toward design of combination therapies with improved efficacy.

Our study was in part limited by the impact of cohort composition on program identification using unsupervised deconvolution. The cNMF programs identified here represented the most coherently co-varying signals in the data, and as such, inclusion of more samples or of distinct biological conditions would likely increase the number of meaningful programs. For instance, inclusion of xenografts treated with standard or rational therapies should reveal contextual tumor and TME adaptations to therapy. Understanding these relationships will be pivotal in designing and preclinical testing of more effective rational and combination therapies.

## SUPPLEMENTARY FIGURES

**Supplementary Figure 1.**
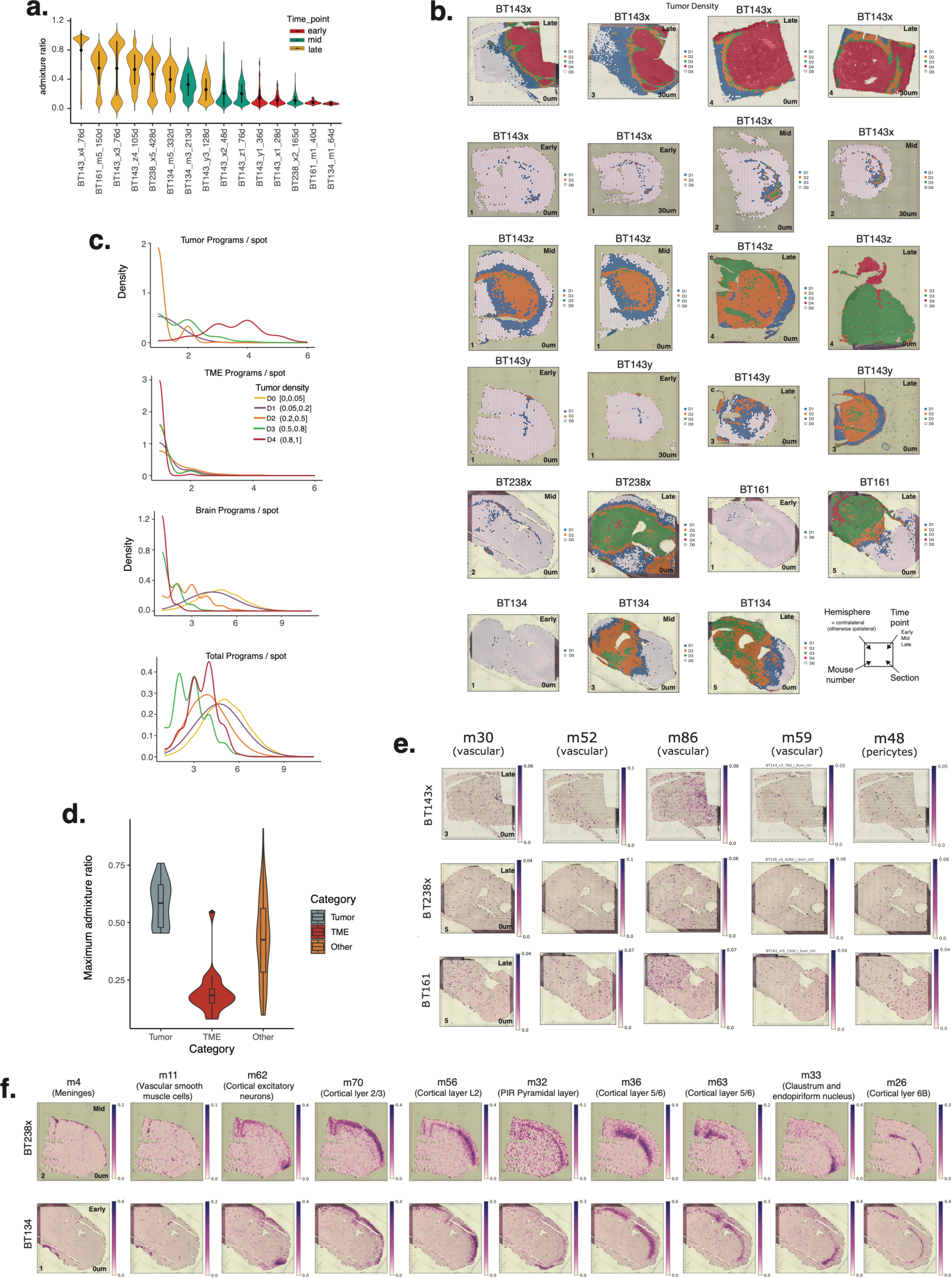
Study overview. **(a)** Distribution of human-mouse admixture per line and time-point (biological replicates merged). Violins are colored based on time-point and ordered by the mean admixture values. **(b)** Spatial plots overlayed with tumor cell density groups (D0-D4) based on human-mouse admixture. **(c)** Density plots of the number of tumor, TME, brain, and total (human and mouse) programs used per spot across tumor cell density groups, D0-D4. A usage threshold of 0.05 was set to define is a program was observed in each spot. **(d)** Violin plots indicate the range of usage values in tumor (n=15), TME(n=19) or other programs (n=81). **(e-f)** Spatial plots of program usage in **(e)** vascular, and **(f)** cortical layer programs.

**Supplementary Figure 2.**
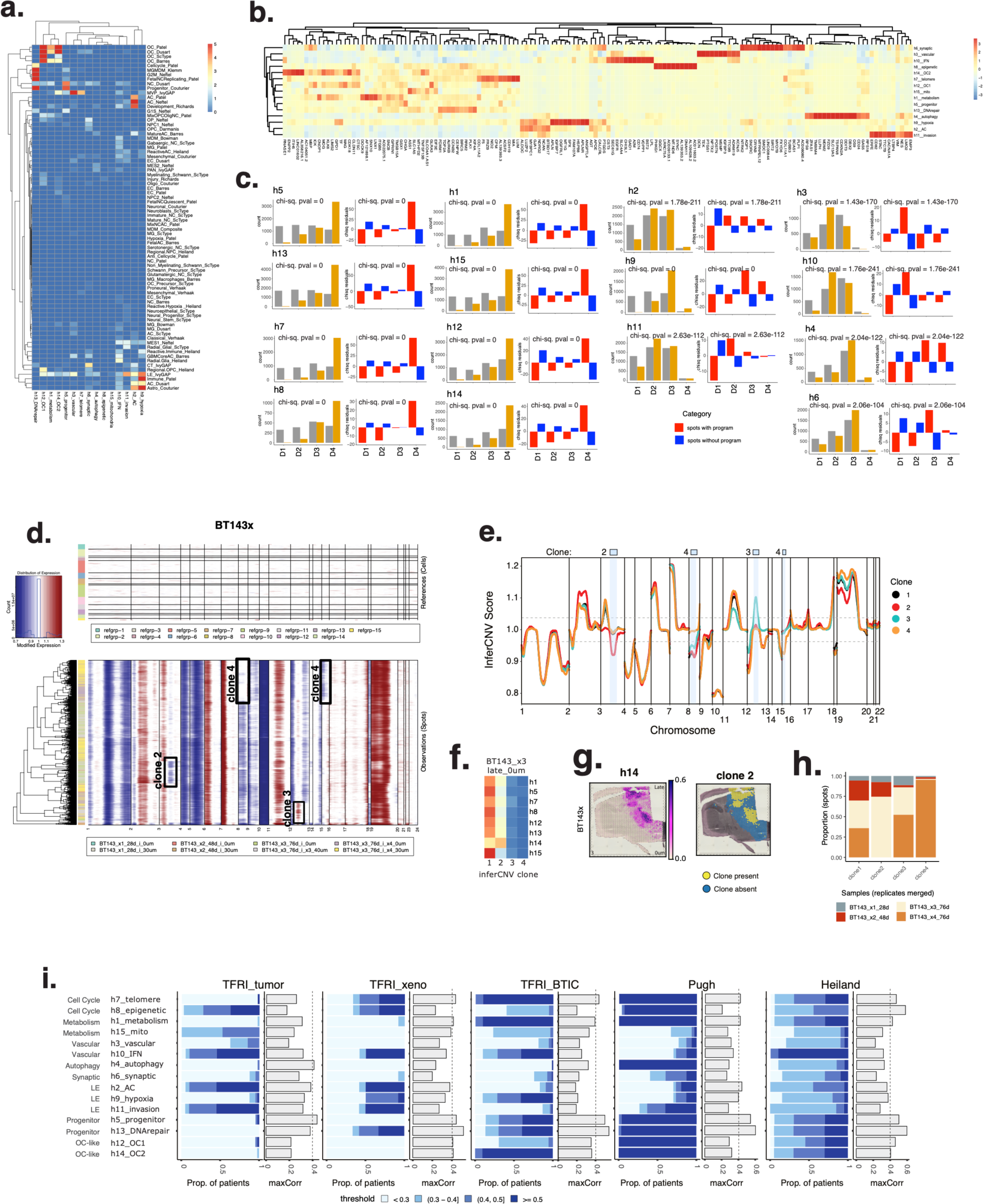
Tumor programs. **(a)** Heatmap of marker-gene scores of tumor programs (columns) calculated with gene sets of GBM cell-states (rows) in external cohorts. **(b)** A hierarchically clustered heatmap of 10 highest-ranking genes in each program, based on cNMF gene scores. **(c)** Number of expected and observed spots with usage of selected programs, and chi-squared residuals and pvalues indicating the significance of difference between observed and expected numbers across categories (D1-D4). **(d)** inferCNV results for BT143x samples with clonal CNVs marked in black outline and labelled. **(e)** Smoothed inferCNV signal for the 4 clones in panel d, with regions of clonal copy number divergence highlighted in blue and labelled above the plot. Divergence is relative to the baseline clone 1, and above/below the score thresholds of 1.5 and 0.5. **(f)** Heatmap of spatial overlap between CNV clones and tumor programswith high usage in BT143×3 and BT143×4 samples. **(g)** Spatial plots of program usage in h14 and clone2 spots in BT143×3. **(i)** Prevalence (blue stacked barplots) and correlation (grey bars) to programs identified in external cohorts. Barplots show the maximum correlation obtained between each tumor program in the xenograft cohort programs derived from cNMF across a series of ranks in external cohorts. A vertical line is drawn at r=0.4 as a visual reference. Prevalence barplots represent the proportion of patients in the external dataset that have usage of the program with maximum correlation above a specific threshold.

**Supplementary Figure 3.**
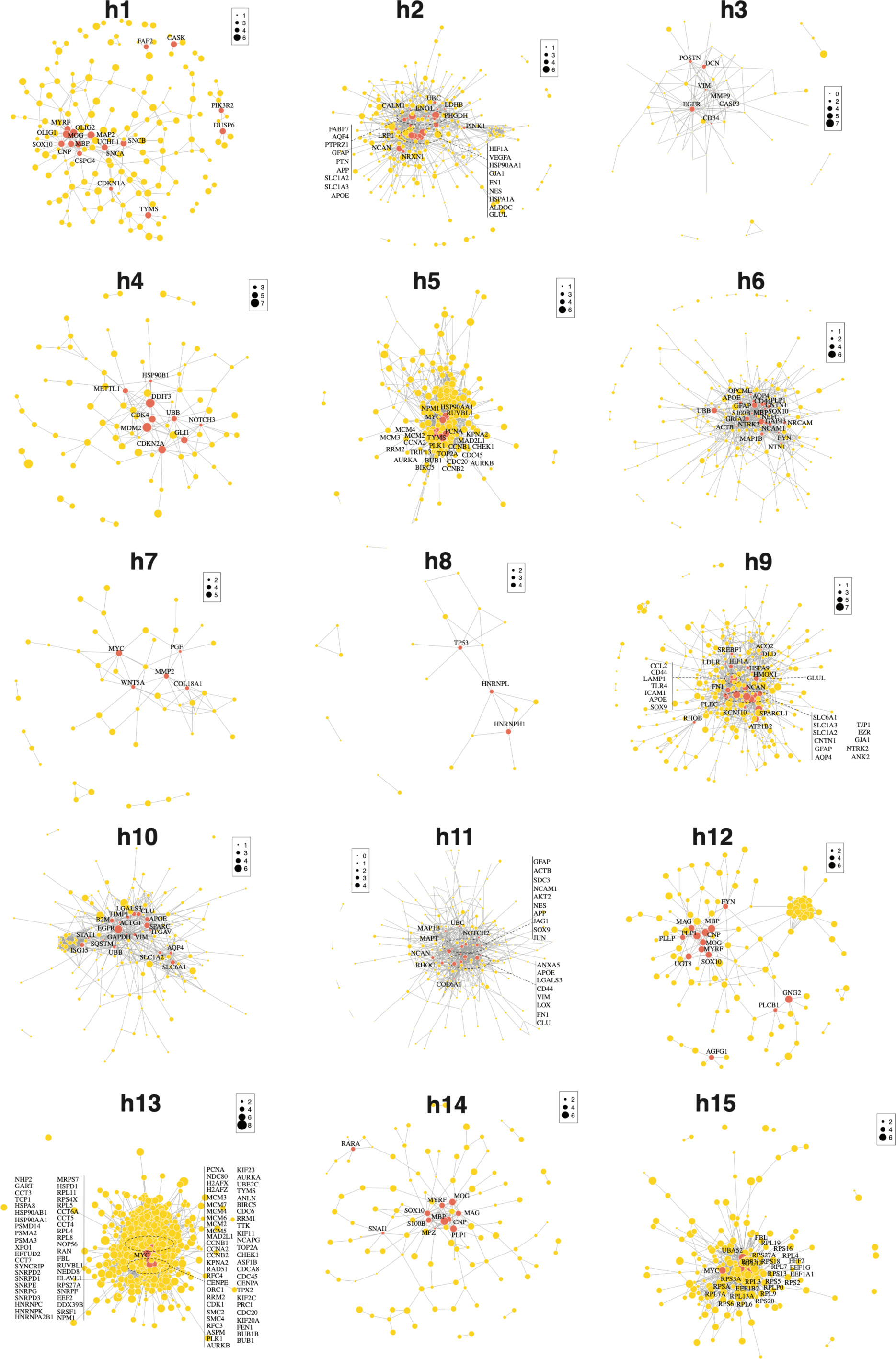
Survival associations. **(a)** Network plot of top-scoring genes (yellow nodes) for tumor programs, with hub genes in orange, and labelled with gene name. Node size is proportional to the correlation between a gene’s usage in a program to its expression, and edges represent the functional associations.

**Supplementary Figure 4.**
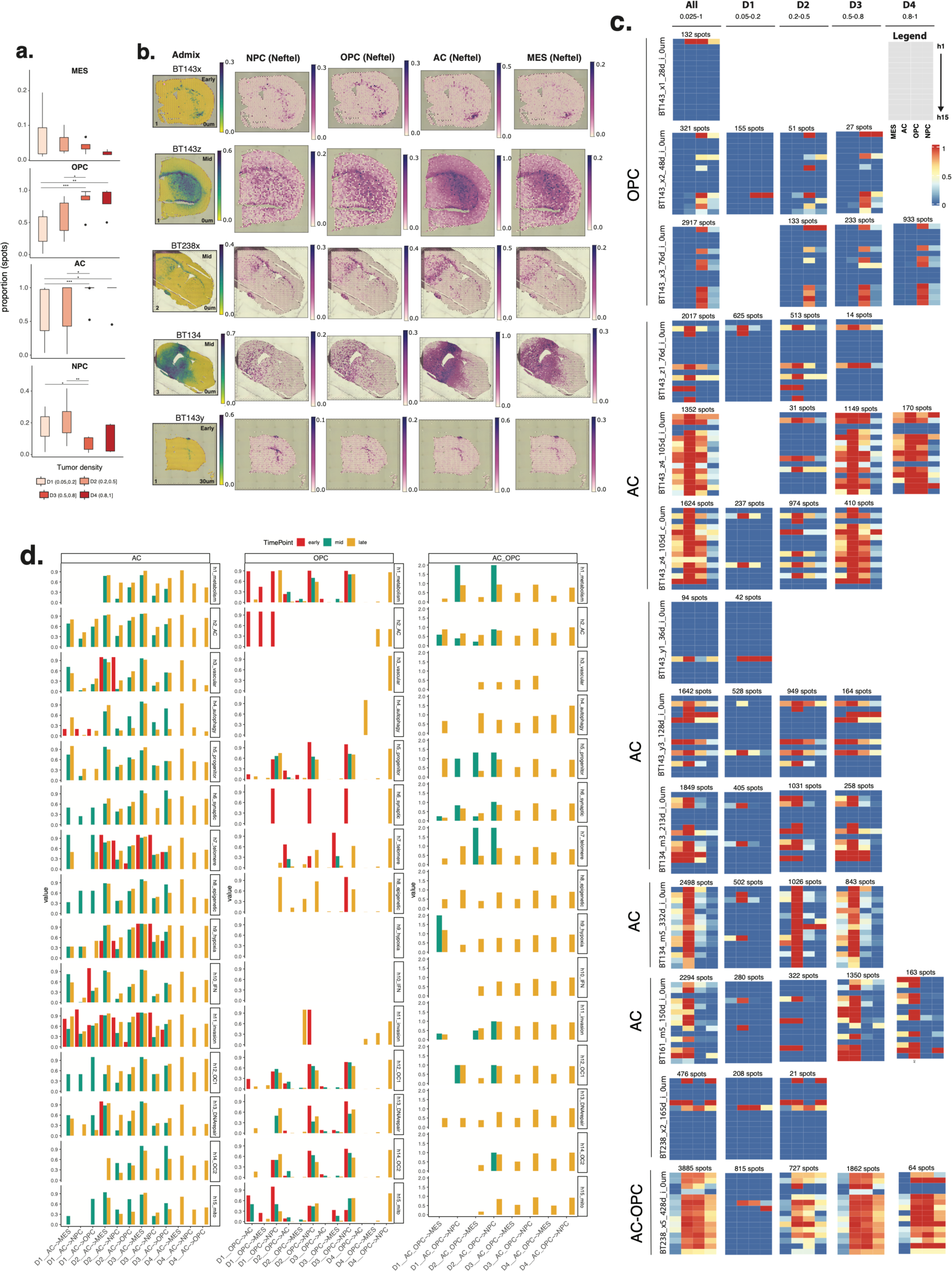
Neftel and baseline states. **(a)** Boxplot showing the proportion of spots in each GBM cell-state with increasing tumor density, D1 to D4. Significant differences across the four groups in each GBM cell-state are marked (wilcoxon rank-sum test ; ns: p > 0.05; *: p <=0.05; **: p <= 0.01). **(b)** Spatial plots of tumor admixture (left) and module scores for NPC, OPC, AC and MES-like states (right). **(c)** Heatmaps depicting the proportion of spatial co-occurence between tumor programs (rows) and GBM cellular states (NPC, OPC, AC and MES; columns) stratified by tumor stage (early, mid, late) and tumor density (D1-D4). Tumor baseline states (labelled) correspond to the most abundant GBM cellular state across all samples from each tumor. **(d)** Barplots indicate proportion of cell-state transitions away from the baseline states (AC, OPC, AC-OPC) for tumor programs, stratified by tumor stage and density. Cell-state transition indicate the proportion of spots with each non-baseline state relative to spots with the baseline state (for spots with cell state usage > 0.1).

**Supplementary Figure 5.**
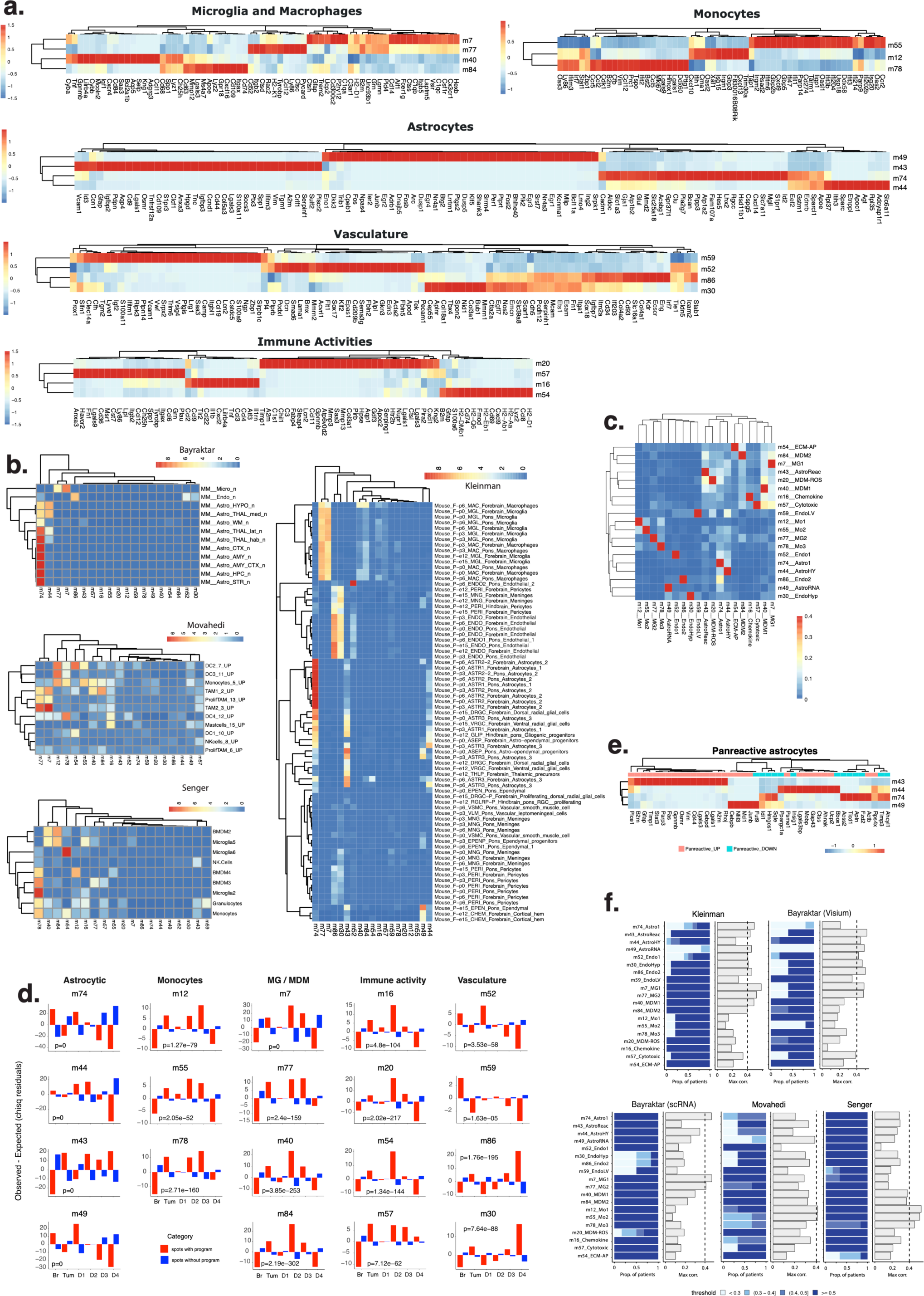
TME programs. **(a)** Scaled heatmaps of the top 20 highest scoring genes in TME programs. **(b)** Heatmaps of marker gene enrichment scores for each TME program (columns) calculated with gene sets of cell-types (rows) identified in external cohorts. **(c)** Proportion of spatial overlap between TME programs. Spatial overlap for a pair of programs is calculated as the proportion of spots in program 1 that also have usage of program 2, and vice versa. **(d)** Chi-squared residuals indicating the observed minus expected number of spots with usage of a specific program per category. Categories include individual tumor density ranges (D1-D4), all tumor regions together (D1-D4 merged), and non-tumor regions (D0; Brain) (x-axis). Chi-sq pvalues indicate the significance of the observed versus expected program usage across categories. **(e)** Scaled heatmap of gene ranks for pan-reactive astrocyte markers indicating the relative contribution of marker genes to program identity. **(f)** Prevalence (blue stacked barplots) and correlation (grey bars) to programs identified in external cohorts. Barplots show the maximum correlation obtained between each TME program in the xenograft cohort programs derived from cNMF across a series of ranks in external cohorts. A vertical line is drawn at r=0.4 as a visual reference. Prevalence barplots represent the proportion of samples in the external dataset that have usage of the program with maximum correlation above a specific threshold.

**Supplementary Figure 6.**
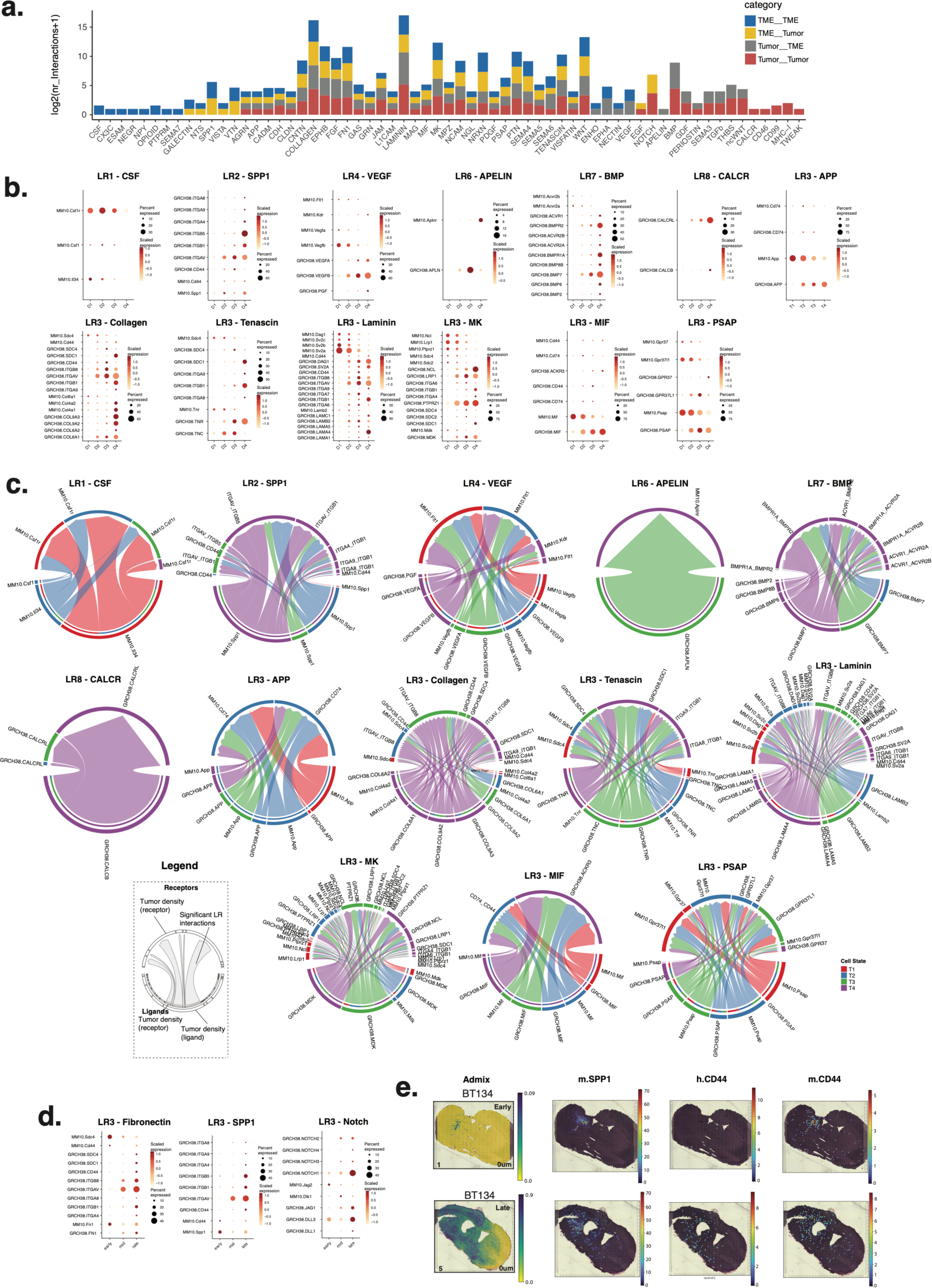
LR communication. **(a)** Stacked barplot indicating the number of interactions per pathway identified as significant. LR interactions are categorized and colored based on the species of the ligand and receptor involved, resulting in 4 interaction categories (ligand receptor): TME TME, TME Tumor, Tumor TME, and Tumor Tumor. **(b)** Scaled expression levels (z-scores) of ligand and receptor genes for selected pathway, stratified by tumor density (D1-D4). **(c)** Chord diagrams show receptor (bottom) and ligand (top) interactions for selected pathways. Interactions are analyzed between all possible pairs of tumor-density groups which make up the sources (outer, lower semi-circle) and targets (upper semi-circle), with links representing the signaling strength of interactions between them (i.e. communication probability). **(d)** Scaled expression levels of ligand and receptor genes for selected pathway, stratified by tumor timepoint (early, mid, late). **(e)** Spatial plots of selected genes (right panels) with tumor admixture for reference (left panels).

**Supplementary Figure 7.**
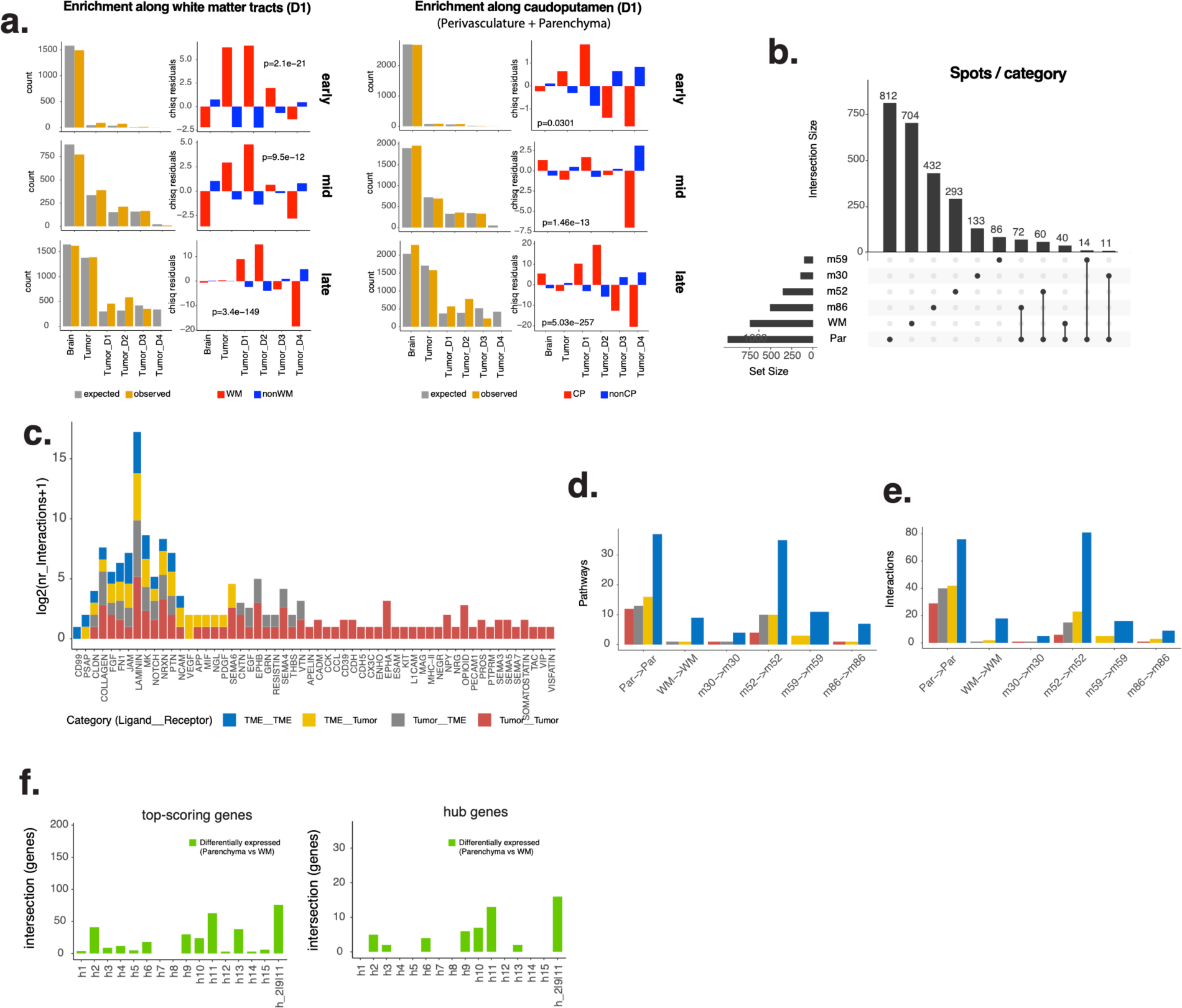
Invasion. **(a)** Number of expected and observed spots with usage of D1 white matter tracts (left panels) and D1 caudoputamen regions (right panels), and chi-squared residuals and pvalues indicating the significance of difference between observed and expected numbers across categories (D0 (Brain), D1-D4 (Tumor), D1-D4 (each)). **(b)** UpSet plot indicating overlap between D1 spot categories. **(c)** Stacked barplot of interactions in significant D1 region pathways. **(d-e)**. Barplots of pathways and interactions within each D1 region of interest. **(f)** Barplot of overlap between top-scoring (left panel) or hub genes (right panel) of each tumor program and the differentially expressed genes that are high in Parenchyma.

## Description of Additional Supplementary Files

### File Name: Supplementary Tables

1. Patient / Cell-line characteristics Description: 1a) Genetic aberrations and clinical data associated with patient-derived cell-lines.
2. Program annotation and gene markers Description: 2a) Gene lists used for annotating human and mouse (normal brain & TME) programs, using *marker gene scores*. 2b) Cell-type / cell-activity / brain structure annotations of mouse and human programs. 2c) Top 1000 genes per tumor program, selected by ranking based on cNMF derived gene-scores. 2d) Top 1000 genes per mouse program.
3. Data associated with tumor programs (Fig. 2) Description: 3a) Pathway enrichment analysis results obtained by querying top 1000 genes in tumor programs against gene-ontology terms in gProfiler. 3b) Marker gene scores for gene-sets representative of GBM cell-types and cell-states, in the tumor programs. 3c) Regulon activity scores per tumor program from SCENIC analysis. 3d) Spatial concordance matrix i.e., short-range spatial overlap between tumor programs (Related to Fig. 2c). 3e) Spatial concordance matrix between tumor and TME programs. 3f) Data and results for the overrepresentation analysis (Chi-sq. test), assessing enrichment of tumor programs across tumor-density groups at the cohort level and 3g) sample level. 3h) Correlation and prevalence of tumor programs in external patient cohorts (Related to Supplementary Fig. 2i).
4. Data associated with survival analyses in tumor programs (Fig. 3) Description: 4a) List of top-scoring genes and 4b) hub genes in tumor programs. 4c) List of top-scoring genes and 4d) hub genes in TME programs. 4e) Summary of survival analyses performed with the TCGA-GBM cohort using top-scoring/hub genes in tumor programs (Tables related to Fig. 3c,f,i). 4f) GSEA results for tumor program top-scoring/hub gene-sets in the TCGA-GBM cohort, used to rank patients in the survival analysis. 4g) Correlation between GSEA-NES of tumor programs.
5. Data associated with TME/Mouse programs Fig. 4) Description: 5a) Pathway enrichment analysis results obtained by querying top 1000 genes in TME programs against gene-ontology terms in gProfiler. 5b) Marker gene scores for gene-sets representative of TME cell-types and cell-states, in the tumor programs. 5c) Regulon activity scores per mouse (brain and TME) program from SCENIC analysis. 5d) Spatial concordance matrix i.e., short-range spatial overlap between tumor programs (Related to Supplementary Fig. 5c). 5e) Data and results for the overrepresentation analysis (Chi-sq. test), assessing enrichment of TME programs across tumor-density groups at the cohort level. 5f) Correlation and prevalence of TME programs in external patient cohorts (Related to Supplementary Fig. 5f).
6. Data associated with Ligand-Receptor analyses across D1 – D4 groups (Fig. 5) Description: 6a) Significant inter and intra-species ligand-receptor interactions identified within and between TME and tumor, across tumor density groups (D1-D4) using Cellchat. In addition to interaction metrics provided by the tool, the table also includes program annotations for ligand and receptor genes. 6b) Number of interactions per pathway in each ligand_receptor category, stratified by tumor density groups (Related to Fig. 5a). 6c) Total number of L-R pairs per pathway in each ligand_receptor category (Related to Fig. 5b). 6d) Pathway enrichment analysis results obtained by querying ligand and receptor genes from the CellChat analysis against gene-ontology terms in gProfiler.
7. Data associated with invasion route programs Description: 7a) Data and results for the overrepresentation analysis (Chi-sq. test), assessing enrichment of white matter and parenchyma programs across tumor-density groups at the cohort level. 7b) Significant inter and intra-species ligand-receptor interactions identified within and between TME and tumor, across invasion-associated programs in D1 spots using Cellchat. 7c) Differential expression analysis results of high tumor density vs. low tumor density spots, from ALDEx2 R package. 7d) Differential expression analysis results of white matter vs. Parencyhma (vascular and parenchymal routes) performed per sample, using Seurat. 7e) Differential expression analysis results of white matter vs. Parenchyma performed per cell-line, using Seurat.

## METHODS

### Intracranial BTIC Models

Xenograft samples were generated from brain tumor initiating cells (BTICs) established from patients with primary and recurrent GBM_[DS1]_ that were maintained as previously described_[DS2]_ prior to intracranial implantation into 6 to 8-week-old female SCID mice_[DS3]_. Some BTIC lines were generated from patient surgical tissue collected from the center of the tumour (x), the highly vascular contrast-enhancing regions (y) or the infiltrating edge (z) (BT143x, BT143y, BT143z, BT238x, BT238z). Mice were housed in groups of three to five and maintained on a 12-hr light/dark schedule with a temperature of 22 °C±1 °C and a relative humidity of 50 ± 5% and provided food and water ad libitum. All animal procedures were reviewed and approved by the University of Calgary Animal Care Committee (Animal Protocol #AC22-0053).

### Visium library preparation and sequencing

The brain was removed and dissected into 2 mm coronal slabs. Fresh tissues were embedded in Tissue Tek OCT compound (Fisher Scientific 14-373-65) and snap-frozen in a chilled isopentane and dry ice bath. 10 µm cryosections were mounted on barcoded Visium slides (10x Genomics), and libraries prepared with the 10x Visium Spatial Gene Expression kit per manufacturer’s protocol. Briefly, sections were fixed in methanol, stained with H&E, and scanned on an EVOS FL Auto Imaging System (Thermo Scientific) using a 10x objective. Permeabilization, reverse transcription, second strand synthesis, denaturation and cDNA synthesis were performed as per protocol. Cycle number determination for cDNA amplification was performed on a BioRad Realtime qPCR system using KAPA SYBR FAST qPCR Master Mix (Roche, KK4600). cDNA QC and qualification was performed on an Agilent 2100 Bioanalyzer with Agilent High sensitivity DNA chips (Agilent 5067-4626). After enzymatic fragmentation and double size selection using SPRIselect reagent (Beckman Coulter, B23318), unique indexes and P5 and P7 Illumina primers were added to the libraries using Dual Index Kit TT Set A (PN-1000215). Libraries were sequenced on an Illumina NextSeq500/550 instrument using paired-end sequencing with the following parameters: 28 cycles for Read1 and 90 cycles for Read, 10-10 cycles for index i7 and index i5, loading concentration 1.8pM on NextSeq 500/550 High Output Kit v2.5 150cycle (Illumina, 20024904).

### Spatial transcriptomics data preprocessing

Illumina sequencing base call data (BCL) was converted to FASTQ files using bcl2fastq (SpaceRanger v1.3.1). Using 10x Genomics SpaceRanger software (v1.3.1), the resulting FASTQ files were mapped to a hybrid genome refence sequence (GRCh38—mm10-2020-A) created by combining the human reference genome (GENCODE v32/Ensembl 98) and mouse reference (GENCODE vM23/Ensembl 98) genome. Data was aligned with STAR v2.7.2a, and mapped to spatial coordinates using the spatial barcode information in SpaceRanger (default parameters). Samples were aggregated using SpaceRanger Aggr. All reads outside the tissue region were removed in the SpaceRanger pipeline. The resulting filtered matrix output is used for subsequent analysis. This matrix consisted of human and mouse genes from 23 samples. The R package Seurat (v4.0.0)^49^ was used to further process this, removing genes with expression in less than 3 spots. Spots with low number of detected genes (< 200) were excluded. Expression of human and mouse genes were normalized together by dividing expression in each spot by the total number of transcripts and multiplied by 10,000, followed by a natural-log transformation.

### Publicly available scRNA-seq and stRNA-seq datasets

Multiple external datasets were downloaded and factorized to yield comparable gene expression programs based on cNMF. The Kleinman^32^ scRNAseq data included 5 normal forebrains (developmental timepoints E12.5, E15.5, P0, P3, and P6), the hindbrain (E12.5), and pons (E15.5, P0, P3, P6). Data was downloaded as CellRanger outputs and cell annotations (GSE133531). Cells in each sample were filtered based on quality control metrics previously described, using the R package Seurat (v4.0.0). Libraries were scaled to 10,000 UMIs per cell and natural log normalized (Seurat::NormalizeData). The Bayraktar^28^ data included matched snRNA-seq and stRNA-seq from adjacent brain sections from 6 adult mice. Data were downloaded from ArrayExpression, including Visium stRNA-seq (E-MTAB-1114; Space Ranger output) and snRNAseq (E-MATB-11115; Cell Ranger output and cell annotations). For Visium profiles, genes with expression in less than 3 spots were discarded. snRNA-seq data was subjected to quality control criteria described in the corresponding publication. Data were scaled to 10,000 UMIs per cell and natural log normalized (Seurat::NormalizeData). The Movahedi^34^ study included CD45+ immune cells from orthotopic GL261 tumors in 3 adult mice. CellRanger outputs and cell annotations were downloaded from www.brainimmuneatlas.org. Outlier cells and low abundance genes were removed based on the workflow previously described. Raw counts were then scaled to 10,000 UMIs per cell and natural log normalized (Seurat::NormalizeData) (v4.0.0). The Senger^7^ study included CD45+ immune cells from orthotopic xenograft models established from patient-derived BTICs. Cell Ranger outputs and cell annotations were downloaded (GSE153487). Data was filtered using quality control metrics previously described and normalized, using R package Seurat (v4.0.0). The Pugh^5^ study included scRNA-seq data from glioblastoma stem cells (GSC) (26 patient GSC cultures) and from tumors (7 patients). Raw and normalized gene expression matrices and cell annotations were downloaded from Broad Institute Single-Cell Portal (https://singlecell.broadinstitute.org/single_cell/study/SCP503). The Heiland^17,30^ study included Visium stRNA-seq profiles of 28 samples from 20 patients. Data from each sample was downloaded in the form of SPATA objects, converted to Seurat objects, and processed with guidelines described in the corresponding publication. The TFRI study included bulk RNA-seq data from 56 patients, including 44 tumor samples, 61 BTIC cell-lines, and 13 xenograft samples, including both longitudinal and multiregional samples. Fastq files were downloaded (EGAS00001002709) and aligned (STAR v2.9.9a^63^, using parameters: --runThreadN 16 --outSAMtype BAM SortedByCoordinate -- quantMode TranscriptomeSAM GeneCounts --outFilterType BySJout --outFilterMatchNminOverLread 0 --outFilterScoreMinOverLread 0 --outSAMstrandField intronMotif --twopassMode Basic). Human reference genome (GENCODE v32/Ensembl 98) was used for mapping tumor and BTIC samples, while a hybrid genome reference (GRCh38—mm10-2020-A) was used for xenografts.

### Admixture calculation

We calculated the ratio of UMI counts from human genes relative to the total UMI count per spot, in order to distinguish tumor from non-neoplastic cells in xenografts. This ratio (human-mouse admixture) represented the transcriptional contribution of tumor cells relative to the total transcriptional output in each spot. Admixture was used to categorize spots based on tumor density: high (admixture 80-100%; D4), moderate (50-80%; D3), low (20-50%; D2), and very low (5-20%; D1). Tumor-free mouse brain was comprised of spots with admixture <5% (D0).

### Selection of over-dispersed (OD) genes

To select for features that are informative of identity and/or activity states in latent space, we selected genes significantly over-dispersed across all spots or samples within each dataset. For the newly-generated xenograft stRNA-seq cohort and the reference sc/snRNA-seq, stRNA-seq datasets, we selected genes detected in more than 1% but less than 100% of spots or single cells that passed quality-control thresholds as described above. Next, genes with higher-than-expected expression variance across the data were selected using a general additive model with a basis of 5 and an adjusted p-value cutoff of 1e-4. Importantly, in case of the xenograft models, feature selection was performed individually for mouse and human genes across filtered spots. The calculations were based on the *restrictCorpus()* function from the R package STDeconvolve (v1.7.0)^26^.

### Matrix factorization and rank selection

We implemented consensus Non-negative Matrix Factorization (cNMF v1.4)^31^, a meta-analysis approach of matrix decomposition to independently infer gene expression programs. The command line version of cNMF was run separately for each dataset. In the xenograft models, the pipeline was run separately for mouse and human gene expression data. Run parameters were first prepared by providing the raw count matrix, the pre-computed normalized matrix, and a list of over-dispersed genes to be used for the factorization steps, all in tab-delimited text file formats to *cnmf prepare*. Prior to this step, spots with no expression of the OD genes were excluded. The number of factors (K) were set to range from 2 to 100 factors. The frobenius loss function was used. Other parameters included --n-iter 200, --total-workers 1, --seed 123456, --densify False. Default parameters were used in the next series of steps, *cnmf factorize* and *cnmf combine*, which factorize and merge results from 200 iterations. The final step, *cnmf consensus,* used to obtain consensus estimates of gene expression programs was performed for each value of K.

We selected an optimal rank based on the trade-off between stability and error, as well as biological interpretability of the factors. This included the anticipated number of cell types and states within a dataset, and manual review of spatial program profiles (in case of stRNA-seq), as well as enrichment of previously annotated cell-types in reference scRNA-seq cohorts. For the selected representative rank in each dataset, the program usage matrix was normalized such that usage values per spot / cell sum to 1 for downstream analyses. In the xenograft models, we selected a rank of 15 for human and 90 for mouse (brain + TME). To quantify the signal of human and mouse programs relative to the total signal, human program usages per spot were multiplied by the admixture ratio. Mouse program usages per spot were multiplied by 1-admixture ratio. The admix-scaled usage matrices were used for further analyses.

In reference datasets we did not select an optimal rank, as these were used to assess the maximum concordance of the xenograft programs with programs found in each external cohort. Thus, we calculated all pairwise correlations between each xenograft program and each program (from all ranks) in the reference datasets, to identify a best match (i.e., a reference program with maximum correlation to the xenograft cohort programs). Further, we calculated the proportion of patients in each external cohort that had usage of the maximally-correlated program above a specific threshold (< 0.3, (0.3-0.4], (0.4-0.5] and > 0.5).

### Annotation of Gene Expression Programs

Gene expression programs identified in the xenograft models were annotated based on cell-type and cell-state gene signatures derived from multiple studies of the mouse brain, GBM, and the GBM microenvironment. The sources and the gene markers used are listed in **Supplementary Table 2a**. We used two methods: 1. Calculation of marker gene scores and 2. Pathway enrichment analysis using g:Profiler^64^.

#### Marker gene scores

We computed marker-gene scores for each program, defined as the rank weighted overlap of top 50 genes in the program with the gene-set being queried. Specifically, for a given query gene-set (*g)*, the intersection with the top 50 genes in a program (*p)* was derived. If *n* is the number of overlapping genes (gene_1_, gene_2_, …, gene_n_), the marker gene score MGS*_gp_* is then calculated as:

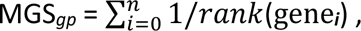

where the rank of a gene is obtained by rank-ordering the genes in each cNMF program using the gene-scores (i.e. the highest scoring gene in a program is rank 1). This method resulted in unambiguous and strong matches to cell-type associated programs, and in poor scores for programs representing a cell-state or activity not matching the reference marker gene sets.

#### Pathway enrichment analysis

To annotate cell activities, we used the top 1000 genes in each program to identify highly enriched pathways from various sources including GO:BP, GO:MF, KEGG and REACTOME, using g:Profiler. Terms with a size > 10 and < 2000 are included and adjusted p-value threshold was set to 0.05. Results were summarized as a heatmap of adjusted p-values of terms ordered by significance in each program.

### Spatial concordance of programs

To infer short-range spatial overlap between program usage among spots for each program, we calculated the proportion of spots that also contribute to a second query program. A usage threshold of >0.1 was needed for a human tumor program to be considered present. Spots with mouse program usage of >0.05 were considered to be positive for that program. The lower threshold used for mouse programs reflects the higher diversity of programs per spot in the normal brain relative to human tumor programs.

### SCENIC

Active regulons in human and mouse programs were identified using the R package SCENIC (v1.1.2.1)^65^ with default parameters. The matrix of z-scores of genes per program obtained from cNMF was used as input to GENIE3 to infer co-expression modules, where each module consisted of a transcription factor (TF) and its predicted targets based on co-expression. Next, using RcisTarget (v1.6.0)^65^, each module (regulon) was pruned to include only targets for which the motif of the TF was enriched. Finally, AUCell (v1.8.0) was used to calculate regulon activity scores per program as the Area Under the Curve (AUC).

### CNV analysis

Copy number changes in individual spots were identified using the R package inferCNV^66^ (v1.3.3). To obtain a robust signal, raw gene counts from each sample were subjected to a spatially aware smoothing method SPCS^67^ using default parameters. As normal reference dataset, we used a cortex section of human brain from 10x Genomics profiled with the Visium platform (https://support.10xgenomics.com/spatial-gene-expression/datasets/1.1.0/V1_Human_Brain_Section_1). The data were then passed to inferCNV for copy number variant inference, separately for each cell line and the following parameters were used: “denoise”, cutoff = 0.1, sd_amplifier = 2.5, scale_data = TRUE, analysis_mode = ’subclusters’, window_length = 201, num_ref_groups = 15. All remaining parameters were set to default. Mitochondrial genes were excluded from this analysis. Individual CNV scores were averaged across clusters to visualize unique transcriptional tumour clones. To further assess how inferred CNVs may impact tumour biology, clones with variable CNVs across chromosome arms were identified by assigning genes to regions of gain or loss using inferCNV score cutoffs of 2.5 times the standard deviation below and above the mean. Spots harbouring these clones were then used to assess spatial concordance with tumor programs.

### GBM cell-state signatures

#### Module score and change across tumor density

Gene signatures for GBM cell-states (astrocyte-like (AC-like), oligodendrocyte-like (OPC-like), neural progenitor-like 1/2 (NPC-like) and mesenchymal-like 1/2 (MES-like)) were obtained from Neftel et al^4^. Spots with admixture ratio of >= 0.05 were selected for analysis of tumor-cell states. Spots were scored for cell-states using the AddModuleScore() function in Seurat (v4.0.0). NPC1/2 and MES1/2 scores were averaged to represent a score for NPC-like and MES-like states respectively. To analyze changes in GBM cell-states as the density of tumor cells in the xenograft models increase (D1-D4), each spot was assigned to the cell-state with the highest module score. Then significance of changes in proportion of spots belonging to each state across the 4 groups were assessed using a Wilcoxon signed rank-sum test and visualized as boxplots.

#### Cell-state transition

To deduce plasticity among GBM cell-states for each human transcriptional program, we calculated a transition score representing change from the baseline of the sample to an end state observed in a given spot. First, a baseline-state was established per sample. This was done by calculating the proportional overlap of highly contributing spots in a program (usage > 0.1) with spots assigned a GBM cell-state based on their highest module score. The overlap metrics are assessed per sample and further by tumor-density groups. The baseline state of a sample was then defined as that which is observed as the majority signal in end-point sample of patient (heatmaps and manual review). Next, the transition score for spots of a given baseline-state at a given time-point (early, mid, or late) and tumor-density range (D1-D4) was derived in two steps: 1. A matrix of proportional overlap between tumor programs and GBM cell-states was created as described above. 2. For each program, the transition score was calculated as the relative contribution of a cell-state over the baseline state and visualized as barplots, grouped by tumor stage and tumor density.

### Cell-cell communication

Inference of ligand-receptor (LR) communication was based on the R package CellChat (v1.6.1)^45^. Given the expected cross-species communication in the xenograft models, ligand, and receptor information available in species-specific CellChatDB were used to create two additional databases, one enumerating human-to-mouse interactions and the other, mouse-to-human. To enable assessment of all directional interactions, the four databases (comprising tumor-to-tumor, TME-to-TME, tumor-to-TME and TME-to-tumor interactions) were re-built into a single database for further analysis. The log-normalized matrix consisting of mouse and human gene expression was used to create a CellChat object. Interactions were inferred across 1. D1-D4 groups and 2. white-matter (WM), caudoputamen (parenchymal), m30, m52, m59 and m86 (vascular) mouse programs only within the D1 group. Spots within D1 (admixture: 0.05-0.2) were assigned to programs in (2) as follows: Spots with usage > 0.1 in m19 or m71 were assigned to WM, > 0.1 in m1 or m73 to parenchymal and > 0.01 in m30, m52, m59 or m86 to the respective vascular programs. In case of spots with more than one of these programs, label assignment between the groups was prioritized as WM > Vascular > Parenchymal. In both analyses, overexpressed genes and interactions were identified using default parameters. Next, *computeCommunProb()* was used to derive significant (p.adj < 0.05) communications with 10% truncated mean for calculating average gene expression per spot. A data-frame of all communications inferred were obtained using *subsetCommunication()* and used for downstream analysis. Chord diagrams of LR interactions per pathway were visualized using *netVisual_chord_cell()*. In (1), pathways were categorized into 8 groups LR1-LR8 based on combinations of the 4 interaction categories (TME TME, TME Tumor, Tumor TME, and Tumor Tumor). Pathway enrichment analysis with GO:BP was performed by querying ligands and receptors of significant interactions in all pathways using g:Profiler. Only terms with a size > 10 and < 2000 were included and adjusted p-value threshold was set to 0.05. Results were summarized as a heatmap of adjusted p-values of terms ordered by significance in each LR group.

### STRINGdb networks and selection of hub genes

#### Selection of hub genes

To build a network of protein-protein interactions per tumor program, we queried a set of “top-scoring” genes in STRING database (v2.6.5)^68^ to obtain known and predicted interactions. Top-scoring genes of a program were derived by plotting the distribution of its gene-scores produced by cNMF, followed by selection of genes that comprise the final component of gaussian-mixture model clusters. This was performed with default parameters of *densityMclust()* from the R package mclust (v6.0.0). Known+Predicted interactions obtained medium confidence levels from STRINGdb were plotted as a network with a fruchterman.reingold layout using the R package iGraph (v1.4.2). Nodes represented genes, and node size corresponded to correlation between expression of the gene and GEP usage across all spots. Finally, hub genes within the network were defined as nodes with a PageRank score in the 95^th^ percentile. Hub genes selected using the PageRank algorithm were observed to be distributed over multiple “fast-greedy” communities within the networks.

#### Hub gene characterization

To functionally characterize hub genes identified in PPI networks of highly-scoring genes, we performed an enrichment analysis of GO:BP terms. Enrichment analysis was performed using the R package gProfiler2 (v0.2.2). The hub genes were queried using gost() with default parameters and convert to an enrichResult using the R package enrichplot (v1.14.2). For each program, terms with size > 5 and < 500 and p.adjust < 0.05 were selected and reduced to ∼5 parent terms using iterative thresholds within *go_reduce()* from the rutils package (v0.99.2). The data was then visualized as a dot-plot with the dot size representing the number of terms reduced per parent term.

### Survival analysis

RNA-seq and clinical data for the TCGA-GBM cohort^69^ were downloaded from http://xena.ucsc.edu. For a given sample, normalized enrichment score (NES) for the gene-set comprising hub genes of each tumor program was calculated using the R package fgsea (v1.20.0) with default parameters. Patients were stratified into three groups based on overall survival time – top and bottom 25%, 33% and 50% and Kaplan-Meier curves were plotted for each group using R package survminer (v0.4.9) with, p-values determined by log-rank test. As a complementary method, a Student’s t.test was also performed between NES of the stratified patients. Kaplan-Meier curves for individual genes were obtained from GEPIA2^70^.

### Over-representation analysis

To assess enrichment of a program across regions of varying tumor cell density, Pearson’s chi-square test was performed using a contingency table encompassing the number spots expressing a program, (selected based on their usage) and the remainder of the spots in each category [Brain(D0), Tumor (D1-D4), and four tumor density regions (D1, D2, D3 and D4)]. “Brain” and “Tumor” categories were excluded while testing for enrichment of tumor programs. A usage threshold of >0.05 was used to select spots in each TME programs and >0.1 for tumor and normal-brain programs. The chi-squared residuals indicating the observed minus expected number of spots with usage of a specific program per category (Brain(D0), Tumor (D1-D2), D1, D2, D3 and D4) are then plotted.

### Differential expression analysis

To determine differentially expressed genes between regions of low and high tumor density with a program-agnostic approach, we used the R package ALDEx2 (v1.24.0)^50^. First, spots in each sample were categorized as low and high density (“lowHs” and “higHs”) by selecting those in the 15^th^ and 75^th^ percentile of the sample’s admixture values. Then, the ALDEx2 pipeline was implemented per sample using the *aldex()* function with the following parameters: mc.samples = 256, test = “t”, effect = TRUE, denom = “lvha”, paired.test = FALSE. Genes with BH adjusted p.value < 0.05 and absolute effect size > 1 were selected as differentially expressed in each category. Further, to obtain differentially expressed genes between WM and other groups (Par, m52, m30, m59, m86) within the D1 region, we used *FindAllMarkers()* with default parameters from R package Seurat (v4.0.0), following filtering (min.cells = 3, min.features = 5, customized nFeature_RNA and nCount_RNA filters per sample) and normalization with NormalizeData() of the selected spots. Genes with adjusted p.value < 0.05 and logfc.threshold > 0.25 were selected as differentially expressed.

## Author Contributions

Varsha Thoppey Manoharan: Conceptualization, Data curation, Software, Formal analysis, Validation, Investigation, Visualization, Methodology, Writing – original draft, Writing – review and editing. Aly Abdelkareem: Data curation, Formal analysis, Methodology. Samuel Brown: Data curation, Methodology, Investigation, Visualization. Aaron Gillmor: Data curation, Formal analysis. Courtney Hall: Formal analysis. Heewon Seo: Formal analysis. Kiran Narta: Data curation, Formal analysis. Sean Grewal: Methodology. Ngoc Ha Dang: Resources, Methodology. Bo Young Ahn: Resources, Methodology. Kata Otz: Resources, Methodology. Xueching Lun: Resources, Methodology. Laura Mah: Resources, Methodology, Investigation, Visualization. Franz Zemp: Resources, Supervision, Methodology. Douglas Mahoney: Resources, Supervision, Methodology. Donna Senger: Conceptualization, Resources, Supervision, Methodology, Writing – review and editing. Jennifer Chan: Conceptualization, Resources, Supervision, Methodology, Writing – review and editing. Sorana Morrissy: Conceptualization, Resources, Supervision, Funding Acquisition, Visualization, Methodology, Writing – original draft, Writing – review and editing.

## Data Availability

Space ranger and cNMF processed files of spatial transcriptomics data generated in this study have been deposited at Datadryad (https://doi.org/10.5061/dryad.wpzgmsbv6). The following publicly available data were obtained as follows: (1) scRNA-seq data of developing mice brain were obtained from GSE133531. (2) snRNA-seq and stRNA-seq data of adult mice brain were obtained from ArrayExpression under accession numbers E-MTAB-1114 and E-MATB-11115. (3) scRNA-seq data from CD45+ immune cells from orthotopic tumors were obtained from www.brainimmuneatlas.org. (4) scRNA-seq data of CD45+ immune cells from IL33+ and IL33-orthotopic xenograft models were obtained from GSE153487. (5) scRNA-seq data from glioblastoma stem cells (GSC) and GBM tumor cells were obtained Broad Institute Single-Cell Portal https://singlecell.broadinstitute.org/single_cell/study/SCP503. (6) stRNA-seq data of GBM patient samples were obtained from https://themilolab.github.io/SPATA2/articles/spata-v2-spata-data.html. (7) Raw RNA-seq data of GBM tumors, cell lines and xenografts were obtained from EGA under the accession number EGAS00001002709. (8) TCGA-GBM RNA-seq data were obtained from http://xena.ucsc.edu. Source data underlying main figures are included the Supplementary Tables file. All remaining relevant data are available in the article, Supplementary Information, or from the corresponding authors upon reasonable request.

## Code availability

Code used to perform analyses presented in this manuscript are available on Github at https://github.com/MorrissyLab/GBM_Xenograft_SpatialTranscriptomics.

## REFERENCES

1. Stupp R, Mason WP, van den Bent MJ, Weller M, Fisher B, Taphoorn MJB, et al. Radiotherapy plus Concomitant and Adjuvant Temozolomide for Glioblastoma. New England Journal of Medicine. 2005;352(10).

2. Cuddapah VA, Robel S, Watkins S, Sontheimer H. A neurocentric perspective on glioma invasion. Vol. 15, Nature Reviews Neuroscience. 2014.

3. Wang L, Jung J, Babikir H, Shamardani K, Jain S, Feng X, et al. A single-cell atlas of glioblastoma evolution under therapy reveals cell-intrinsic and cell-extrinsic therapeutic targets. Nat Cancer. 2022;3(12).

4. Neftel C, Laffy J, Filbin MG, Hara T, Shore ME, Rahme GJ, et al. An Integrative Model of Cellular States, Plasticity, and Genetics for Glioblastoma. Cell. 2019;178(4).

5. Richards LM, Whitley OKN, MacLeod G, Cavalli FMG, Coutinho FJ, Jaramillo JE, et al. Gradient of Developmental and Injury Response transcriptional states defines functional vulnerabilities underpinning glioblastoma heterogeneity. Nat Cancer. 2021;2(2).

6. Varn FS, Johnson KC, Martinek J, Huse JT, Nasrallah MP, Wesseling P, et al. Glioma progression is shaped by genetic evolution and microenvironment interactions. Cell. 2022;185(12).

7. De Boeck A, Ahn BY, D’Mello C, Lun X, Menon S V., Alshehri MM, et al. Glioma-derived IL-33 orchestrates an inflammatory brain tumor microenvironment that accelerates glioma progression. Nat Commun. 2020;11(1).

8. Wang J, Xu SL, Duan JJ, Yi L, Guo YF, Shi Y, et al. Invasion of white matter tracts by glioma stem cells is regulated by a NOTCH1–SOX2 positive-feedback loop. Nat Neurosci. 2019;22(1).

9. Jung E, Osswald M, Ratliff M, Dogan H, Xie R, Weil S, et al. Tumor cell plasticity, heterogeneity, and resistance in crucial microenvironmental niches in glioma. Nat Commun. 2021;12(1).

10. Sarkar S, Mirzaei R, Zemp FJ, Wei W, Senger DL, Robbins SM, et al. Activation of NOTCH signaling by Tenascin-C promotes growth of human brain tumor-initiating cells. Cancer Res. 2017;77(12).

11. Comba A, Faisal SM, Dunn PJ, Argento AE, Hollon TC, Al-Holou WN, et al. Spatiotemporal analysis of glioma heterogeneity reveals COL1A1 as an actionable target to disrupt tumor progression. Nat Commun. 2022;13(1).

12. Bastola S, Pavlyukov MS, Yamashita D, Ghosh S, Cho H, Kagaya N, et al. Glioma-initiating cells at tumor edge gain signals from tumor core cells to promote their malignancy. Nat Commun. 2020;11(1).

13. Minata M, Audia A, Shi J, Lu S, Bernstock J, Pavlyukov MS, et al. Phenotypic Plasticity of Invasive Edge Glioma Stem-like Cells in Response to Ionizing Radiation. Cell Rep. 2019;26(7).

14. Venkatesh HS, Tam LT, Woo PJ, Lennon J, Nagaraja S, Gillespie SM, et al. Targeting neuronal activity-regulated neuroligin-3 dependency in high-grade glioma. Nature. 2017;549(7673).

15. Bressan D, Battistoni G, Hannon GJ. The dawn of spatial omics. Vol. 381, Science. 2023.

16. Greenwald AC, Darnell NG, Hoefflin R, Simkin D, Gonzalez-Castro LN, Mount C, et al. Integrative spatial analysis reveals a multi-layered organization of glioblastoma. bioRxiv. 2023;

17. Ravi VM, Will P, Kueckelhaus J, Sun N, Joseph K, Salié H, et al. Spatially resolved multi-omics deciphers bidirectional tumor-host interdependence in glioblastoma. Cancer Cell. 2022;40(6).

18. Zheng Y, Carrillo-Perez F, Pizurica M, Heiland DH, Gevaert O. Spatial cellular architecture predicts prognosis in glioblastoma. Nat Commun. 2023;14(1).

19. Gangoso E, Southgate B, Bradley L, Rus S, Galvez-Cancino F, McGivern N, et al. Glioblastomas acquire myeloid-affiliated transcriptional programs via epigenetic immunoediting to elicit immune evasion. Cell. 2021;184(9).

20. Ren Y, Huang Z, Zhou L, Xiao P, Song J, He P, et al. Spatial transcriptomics reveals niche-specific enrichment and vulnerabilities of radial glial stem-like cells in malignant gliomas. Nat Commun. 2023;14(1).

21. Andersson A, Bergenstråhle J, Asp M, Bergenstråhle L, Jurek A, Fernández Navarro J, et al. Single-cell and spatial transcriptomics enables probabilistic inference of cell type topography. Commun Biol. 2020;3(1).

22. Cable DM, Murray E, Zou LS, Goeva A, Macosko EZ, Chen F, et al. Robust decomposition of cell type mixtures in spatial transcriptomics. Nat Biotechnol. 2022;40(4).

23. Biancalani T, Scalia G, Buffoni L, Avasthi R, Lu Z, Sanger A, et al. Deep learning and alignment of spatially resolved single-cell transcriptomes with Tangram. Nat Methods. 2021;18(11).

24. Ma Y, Zhou X. Spatially informed cell-type deconvolution for spatial transcriptomics. Nat Biotechnol. 2022;40(9).

25. Elosua-Bayes M, Nieto P, Mereu E, Gut I, Heyn H. SPOTlight: Seeded NMF regression to deconvolute spatial transcriptomics spots with single-cell transcriptomes. Nucleic Acids Res. 2021;49(9).

26. Miller BF, Huang F, Atta L, Sahoo A, Fan J. Reference-free cell type deconvolution of multi-cellular pixel-resolution spatially resolved transcriptomics data. Nat Commun. 2022;13(1).

27. Rodriques SG, Stickels RR, Goeva A, Martin CA, Murray E, Vanderburg CR, et al. Slide-seq: A scalable technology for measuring genome-wide expression at high spatial resolution. Science (1979). 2019;363(6434).

28. Kleshchevnikov V, Shmatko A, Dann E, Aivazidis A, King HW, Li T, et al. Cell2location maps fine-grained cell types in spatial transcriptomics. Nat Biotechnol. 2022;40(5).

29. Coleman K, Hu J, Schroeder A, Lee EB, Li M. SpaDecon: cell-type deconvolution in spatial transcriptomics with semi-supervised learning. Commun Biol. 2023;6(1).

30. Shen Y, Grisdale CJ, Islam SA, Bose P, Lever J, Zhao EY, et al. Comprehensive genomic profiling of glioblastoma tumors, BTICs, and xenografts reveals stability and adaptation to growth environments. Proc Natl Acad Sci U S A. 2019;116(38).

31. Kotliar D, Veres A, Nagy MA, Tabrizi S, Hodis E, Melton DA, et al. Identifying gene expression programs of cell-type identity and cellular activity with single-cell RNA-Seq. Elife. 2019;8.

32. Jessa S, Blanchet-Cohen A, Krug B, Vladoiu M, Coutelier M, Faury D, et al. Stalled developmental programs at the root of pediatric brain tumors. Nat Genet. 2019;51(12).

33. Lein ES, Hawrylycz MJ, Ao N, Ayres M, Bensinger A, Bernard A, et al. Genome-wide atlas of gene expression in the adult mouse brain. Nature. 2007;445(7124).

34. Pombo Antunes AR, Scheyltjens I, Lodi F, Messiaen J, Antoranz A, Duerinck J, et al. Single-cell profiling of myeloid cells in glioblastoma across species and disease stage reveals macrophage competition and specialization. Nat Neurosci. 2021;24(4).

35. Couturier CP, Ayyadhury S, Le PU, Nadaf J, Monlong J, Riva G, et al. Single-cell RNA-seq reveals that glioblastoma recapitulates a normal neurodevelopmental hierarchy. Nat Commun. 2020;11(1).

36. Puchalski RB, Shah N, Miller J, Dalley R, Nomura SR, Yoon JG, et al. An anatomic transcriptional atlas of human glioblastoma. Science (1979). 2018;360(6389).

37. Patel AP, Tirosh I, Trombetta JJ, Shalek AK, Gillespie SM, Wakimoto H, et al. Single-cell RNA-seq highlights intratumoral heterogeneity in primary glioblastoma. Science (1979). 2014;344(6190).

38. Dusart P, Hallström BM, Renné T, Odeberg J, Uhlén M, Butler LM. A Systems-Based Map of Human Brain Cell-Type Enriched Genes and Malignancy-Associated Endothelial Changes. Cell Rep. 2019;29(6).

39. Darmanis S, Sloan SA, Croote D, Mignardi M, Chernikova S, Samghababi P, et al. Single-Cell RNA-Seq Analysis of Infiltrating Neoplastic Cells at the Migrating Front of Human Glioblastoma. Cell Rep. 2017;21(5).

40. Verhaak RGW, Hoadley KA, Purdom E, Wang V, Qi Y, Wilkerson MD, et al. Integrated Genomic Analysis Identifies Clinically Relevant Subtypes of Glioblastoma Characterized by Abnormalities in PDGFRA, IDH1, EGFR, and NF1. Cancer Cell. 2010;17(1).

41. Doetsch F, García-Verdugo JM, Alvarez-Buylla A. Cellular composition and three-dimensional organization of the subventricular germinal zone in the adult mammalian brain. Journal of Neuroscience. 1997;17(13).

42. Monteiro AR, Hill R, Pilkington GJ, Madureira PA. The role of hypoxia in glioblastoma invasion. Vol. 6, Cells. 2017.

43. Venkataramani V, Yang Y, Schubert MC, Reyhan E, Tetzlaff SK, Wißmann N, et al. Glioblastoma hijacks neuronal mechanisms for brain invasion. Cell. 2022;185(16).

44. Das S, Li Z, Noori A, Hyman BT, Serrano-Pozo A. Meta-analysis of mouse transcriptomic studies supports a context-dependent astrocyte reaction in acute CNS injury versus neurodegeneration. J Neuroinflammation. 2020;17(1).

45. Jin S, Guerrero-Juarez CF, Zhang L, Chang I, Ramos R, Kuan CH, et al. Inference and analysis of cell-cell communication using CellChat. Nat Commun. 2021;12(1).

46. Angenendt L, Wöste M, Mikesch JH, Arteaga MF, Angenendt A, Sandmann S, et al. Calcitonin receptor-like (CALCRL) is a marker of stemness and an independent predictor of outcome in pediatric AML. Blood Adv. 2021;5(21).

47. Gu S, Shu L, Zhou L, Wang Y, Xue H, Jin L, et al. Interfering with CALCRL expression inhibits glioma proliferation, promotes apoptosis, and predicts prognosis in low-grade gliomas. Ann Transl Med. 2022;10(23).

48. Scherer HJ. Structural development in gliomas. American Journal of Cancer. 1938;34(3).

49. Hao Y, Hao S, Andersen-Nissen E, Mauck WM, Zheng S, Butler A, et al. Integrated analysis of multimodal single-cell data. Cell. 2021;184(13).

50. Fernandes AD, Reid JNS, Macklaim JM, McMurrough TA, Edgell DR, Gloor GB. Unifying the analysis of high-throughput sequencing datasets: Characterizing RNA-seq, 16S rRNA gene sequencing and selective growth experiments by compositional data analysis. Microbiome. 2014;2(1).

51. Venkatesh HS, Johung TB, Caretti V, Noll A, Tang Y, Nagaraja S, et al. Neuronal activity promotes glioma growth through neuroligin-3 secretion. Cell. 2015;161(4).

52. Lustig M, Sakurai T, Grumet M. Nr-CAM promotes neurite outgrowth from peripheral ganglia by a mechanism involving axonin-1 as a neuronal receptor. Dev Biol. 1999;209(2).

53. Sehgal A, Boynton AL, Young RF, Vermeulen SS, Yonemura KS, Kohler EP, et al. Cell adhesion molecule Nr-CAM is over-expressed in human brain tumors. Int J Cancer. 1998;76(4).

54. Schuster A, Klein E, Neirinckx V, Knudsen AM, Fabian C, Hau AC, et al. AN1-type zinc finger protein 3 (ZFAND3) is a transcriptional regulator that drives Glioblastoma invasion. Nat Commun. 2020;11(1).

55. Aiken J, Moore JK, Bates EA. TUBA1A mutations identified in lissencephaly patients dominantly disrupt neuronal migration and impair dynein activity. Hum Mol Genet. 2019;28(8).

56. Belvindrah R, Natarajan K, Shabajee P, Bruel-Jungerman E, Bernard J, Goutierre M, et al. Mutation of the α-tubulin Tuba1a leads to straighter microtubules and perturbs neuronal migration. Journal of Cell Biology. 2017;216(8).

57. Camby I, Belot N, Lefranc F, Sadeghi N, De Launoit Y, Kaltner H, et al. Galectin-1 modulates human glioblastoma cell migration into the brain through modifications to the actin cytoskeleton and levels of expression of small GTPases. J Neuropathol Exp Neurol. 2002;61(7).

58. Van Woensel M, Mathivet T, Wauthoz N, Rosière R, Garg AD, Agostinis P, et al. Sensitization of glioblastoma tumor micro-environment to chemo- and immunotherapy by Galectin-1 intranasal knock-down strategy. Sci Rep. 2017;7(1).

59. Shevchenko V, Arnotskaya N, Pak O, Sharma A, Sharma HS, Khotimchenko Y, et al. Molecular determinants of the interaction between glioblastoma CD133+ cancer stem cells and the extracellular matrix. In: International Review of Neurobiology. 2020.

60. Turtoi A, Blomme A, Bianchi E, Maris P, Vannozzi R, Naccarato AG, et al. Accessibilome of human glioblastoma: Collagen-VI-alpha-1 is a new target and a marker of poor outcome. J Proteome Res. 2014;13(12).

61. Sharma N, Atolagbe OT, Ge Z, Allison JP. LILRB4 suppresses immunity in solid tumors and is a potential target for immunotherapy. Journal of Experimental Medicine. 2021;218(7).

62. Xiong A, Zhang J, Chen Y, Zhang Y, Yang F. Integrated single-cell transcriptomic analyses reveal that GPNMB-high macrophages promote PN-MES transition and impede T cell activation in GBM. EBioMedicine. 2022;83.

63. Dobin A, Davis CA, Schlesinger F, Drenkow J, Zaleski C, Jha S, et al. STAR: Ultrafast universal RNA-seq aligner. Bioinformatics. 2013;29(1).

64. Kolberg L, Raudvere U, Kuzmin I, Adler P, Vilo J, Peterson H. G:Profiler-interoperable web service for functional enrichment analysis and gene identifier mapping (2023 update). Nucleic Acids Res. 2023;51(W1).

65. Aibar S, González-Blas CB, Moerman T, Huynh-Thu VA, Imrichova H, Hulselmans G, et al. SCENIC: Single-cell regulatory network inference and clustering. Nat Methods. 2017;14(11).

66. Tickle T, Tirosh I, Georgescu C, Brown M, Haas B. inferCNV of the Trinity CTAT Project. 2019.

67. Liu Y, Wang T, Duggan B, Sharpnack M, Huang K, Zhang J, et al. SPCS: a spatial and pattern combined smoothing method for spatial transcriptomic expression. Brief Bioinform. 2022;23(3).

68. Szklarczyk D, Franceschini A, Wyder S, Forslund K, Heller D, Huerta-Cepas J, et al. STRING v10: Protein-protein interaction networks, integrated over the tree of life. Nucleic Acids Res. 2015;43(D1).

69. Weinstein JN, Collisson EA, Mills GB, Shaw KRM, Ozenberger BA, Ellrott K, et al. The cancer genome atlas pan-cancer analysis project. Nat Genet. 2013;45(10).

70. Tang Z, Kang B, Li C, Chen T, Zhang Z. GEPIA2: an enhanced web server for large-scale expression profiling and interactive analysis. Nucleic Acids Res. 2019;47(W1).

