## Supplementary_material for "Spatiotemporal modeling reveals high-resolution invasion states in glioblastoma"

### SUPPLEMENTARY MATERIALS

### SUPPLEMENTARY RESULTS

#### Tumor program descriptions

Here we provide additional details regarding pathway annotations for individual programs observed in the human tumor cells.

A first set of six programs represented Progenitor, Cell Cycle, and Metabolic themes. The two Progenitor programs (*h5_progenitor*, *h13_DNArepair*) were distinguished from each-other based on cell state and pathways, with high scores for progenitor and G2M tumor cell states in *h13_DNArepair* (**Fig. 2b, Supplementary Fig. 2a**), and significant enrichment of multiple DNA damage repair terms including base excision and mismatch repair **(Fig. 2a, Supplementary Table 3a)**. In contrast, the *h5_progenitor* program showed higher scores for G1S and neuronal progenitor terms (**Fig. 2b, Supplementary Fig. 2a**). Both programs were present at all timepoints and across density regions, with significantly higher abundance observed in D4, indicating that progenitor states were more prevalent at high tumor cell density (**Fig. 3e, Supplementary Fig. 2c, Supplementary Table 3f**). Next, two Cell Cycle programs (*h7_telomere*, *h8_epigenetic*) were enriched in DNA replication terms. *h7_telomere* was the most prevalent of the two, involving terms for telomere maintenance, and protein localization and degradation; in contrast, major pathway themes in *h8_epigenetic* focused on chromatin biology (e.g. histone modifications via lysine acetylation) **(Fig. 2a, Supplementary Table 3a)**. Finally, we identified two activity programs with clear pathway themes relating to Metabolism – *h1_metabolism* was defined by cholesterol biosynthesis and nucleotide and RNA metabolism, while *h15_mito* terms converged on cellular respiration and mitochondrial translation **(Fig. 2a, Supplementary Table 3a)**. We observed a consistent association of the Progenitor, Cell Cycle, and Metabolic programs with the vascular/astrocytic niche in D4, where these programs had the highest prevalence (**Fig. 2e, Supplementary Table 3e**). A particularly high overlap of *h7_telomere* and *h8_epigenetic* programs with the Progenitor programs and *m30_endoHyp* – indicated that a subset of cycling progenitors was highly concentrated within an aberrant vascular niche in regions of hypoxia (**Supplementary Table 3e**), in line with previous evidence for this stem cell niche[1,2].

A second set of programs represented OC-like, AC-like, and Invasion themes. Two programs (*h12_OC1, h14_OC2*) had strong matches to OC-like cell states (**Fig 2b, Supplementary Fig. 2a**), and were enriched in terms for regulation of oligodendrocyte differentiation **(Fig. 2a, Supplementary Table 3a)**. Assessment of genetic heterogeneity in the stRNAseq data revealed that *h14_OC2* was a genetic subclone present in the BT143x cell line (**Supplementary Fig. 2d-h**), explaining its high similarity to *h12_OC1*, and its nested spatial localization (**Fig. 3c**). Both were most abundant in D4 where they occupied the vascular/astrocyte niche (**Fig. 2e, Supplementary Fig. 2c, Supplementary Table 3e,f**). Next, two AC-like programs (*h2_AC*, *h9_hypoxia*) matched well to AC-like GBM states (**Fig. 2b**, **Supplementary Fig. 2a**), had pathway terms involving hypoxia response/regulation of angiogenesis (*h9_hypoxia*), the GBM developmental state (*h2_AC*), and moderate matches to the leading edge (LE_IvyGAP[3]; both *h2* and *h9*) **(Fig. 2a, Supplementary Table 3a)**. The *h11_invasion* program scored highly for LE_IvyGAP, comprised a majority of D1-D2 spots at early and late timepoints, and was the only tumor program with significant over-representation in D1 regions within individual patients (**Fig**

**2e,g, Supplementary Table 3g**). Together, this trio of programs revealed a gradient of outward expansion, with *h9_hypoxia* most centrally located, *h2_AC* in the surrounding regions, and *h11_invason* in the most outlying parts of the tumor (**Fig. 2e-g**).

While the eleven programs described above were observed in nearly all lines from multiple patients, a final set of four programs was enriched in specific patients (**Fig. 3e**). *h4_autophagy* (BT161) was enriched in pathways involving senescence, autophagy, mitophagy, aggrephagy, and apoptosis **(Fig. 2a, Supplementary Table 3a)**. Additional (lower) matches to mesenchymal and PAN (**Fig 2b**) linked these states to autophagy. In BT238x, we observed high usage of *h6_synaptic* (**Fig. 2d**), with major pathway themes around synapse formation, and cell signaling and adhesion **(Fig. 2a, Supplementary Table 3a)**. Finally, we identified two programs that relate to angiogenesis (BT134). *h3_vascular* had strong matches to microvascular proliferation (**Fig. 2b**), and enrichment in pathways for vascular development, cell migration, and ECM-receptor interactions (**Fig 2a, Supplementary Table 3a**). *h10_IFN* had weaker matches to PAN and mesenchymal states (**Fig 2b**), enrichment of angiogenesis terms, and of interferon signaling terms (**Fig 2a, Supplementary Table 3a**). The observation that these programs were prevalent within individual patients suggested that specific tumor genotypes could be linked to unique repertoires of tumor cell programs, and that larger xenograft cohorts may therefore reveal additional insights. In our cohort we noted high program stability even when including an additional sample, and re-running program identification. The additional sample (BT238z) was a slow-growing and highly diffuse BTIC line established from the invasive front (z) of patient BT238. In this case, we observed nearly identical factorization results for both broadly-used (e.g. *h11_invasion*) and genotype-enriched programs (e.g. *h6_synaptic* specific to BT238) (data not shown).

**Microenvironment program descriptions**

Here we provide additional details regarding pathway annotations for individual programs observed in cells of the tumor microenvironment.

Astrocytic programs included both cell types (*m74_Astro1*, *m44_AstroHY*) and activities (*m43_AstroReac*, *m49_AstroRNA*). *m74_Astro1* was widespread throughout the normal mouse brain, prevalent in the invasive tumor front and excluded from dense tumor regions (**Fig. 4f,h, Supplementary Fig. 5d, Supplementary Table 5e**), and enriched in pathways characteristic of mature astrocytes including metabolic processes (glycolysis, nucleotide metabolism, cellular respiration), regulation of synapse structures and activities, and cellular response to hypoxia and reactive oxygen species (ROS) (**Fig. 4d, Supplementary Table 5a**). Similarly, *m49_AstroRNA* showed broad usage across normal brain with exclusion from dense tumor (**Fig. 4f,h, Supplementary Fig. 5d, Supplementary Table 5e**), likely representing normal astrocytic activities (RNA processing, proliferation, glutamatergic synapses) (**Fig. 4d, Supplementary Table 5a**). In contrast, *m44_AstroHY* localized both to regions of tumor and normal brain (hypothalamus), suggesting it represents a regional astrocyte subtype that can be rapidly recruited to sites of injury, or alternatively, that it is a cell state which astrocytes in the TME can rapidly adopt (**Fig. 4h, Supplementary Fig. 5d, Supplementary Table 5e**). Significant enrichment of terms relating to regulation of precursor cell proliferation highlighted a role for this otherwise normal program in the context of the GBM TME (**Fig. 4d, Supplementary Table 5a**). Next, the *m43_AstroReac* program was the only one enacted across the full spectrum of tumor density, sustained at all timepoints of disease progression, and highly specific to regions of tumor (**Fig. 2f,h, Supplementary Fig. 5d, Supplementary Table 5e**). Co-localization of *m43_AstroReac* with *m74_Astro1* indicated that this was an activity enacted by normal astrocytes in the context of the TME (**Supplementary Fig. 5c**), a conclusion further supported by enrichment of terms relating to proliferation, inflammatory response, and ECM organization, and a strong match to a reactive astrocyte signature[4] (**Fig. 4d, Supplementary Fig. 5e, Supplementary Table 5a**).

We annotated four endothelial programs, including *m52_Endo1*, *m86_Endo2*, *m30_EndoHyp*, and *m59_EndoLV* (**Fig. 4a,c, Supplementary Fig. 1e,5a, Supplementary Table 2b**). Of these, *m52_Endo1* best matched normal endothelial cells in reference datasets, had mature endothelial markers (Gkn3, Sema3g, Alpl, Pecam1; **Supplementary Fig. 5a,b**), and was widespread in the normal brain **(Supplementary Fig. 1e,5d, Supplementary Table 5e)**. *m52_Endo1* had decreased prevalence within regions of high tumor density, where a second endothelial program (*m86_Endo2*) was the major vascular component (**Fig. 2f, Supplementary Fig. 1e,5d, Supplementary Table 5e**). This tumor-enriched program (*m86_Endo2*) was distinguished by pathways relating to endothelial cell migration and proliferation, tight junction organization and assembly, leukocyte activation and cell-cell adhesion, and regulation of coagulation and hemostasis **(Supplementary Fig. 4c, Supplementary Table 5a)**. Together, these terms encompassed aspects of tumor vasculature and blood-brain-barrier remodeling positively associated with increasing tumor density. Another tumor-enriched program (*m30_EndoHyp*) was distinguished by pathways relating to cell proliferation and DNA replication, regulation of cell migration involved in sprouting angiogenesis, collagen fibril organization, and response to hypoxia **(Supplementary Fig. 4c, Supplementary Table 5a)**. This is an activity program sparsely present throughout the brain, but significantly enriched in denser tumor regions with low oxygen availability (**Fig. 2f, Supplementary Fig. 1e,5d, Supplementary Table 5e**). Finally, we identified *m59_EndoLV* as highly scoring for lymph vessel morphogenesis, suggesting the possibility that lymphatic and vascular vessels in GBM are distinguishable with this deconvolution approach. *m59_EndoLV* showed a sparse pattern of usage throughout the brain largely corresponding to known locations of lymphatic vasculature[5] (meninges) and was also present across timepoints and tumor densities (**Fig. 2f, Supplementary Fig. 1e,5d, Supplementary Table 5e**). In support of this program's lymphatic identity, we observed high program scores for lymphangiogenic regulators including Lyve1, Clec14a, Ptpn14, Vwf, and others[5] (**Supplementary Fig. 5f**). Of these, Vwf (the second highest-scoring gene in *m59_EndoLV;* **Supplementary Table 2d**) has a central role in vascular inflammation (Gragnano, Mediators Inflamm 2017), regulation of vessel permeability, and immune cell recruitment[6].

Multiple immune cell types were evident in our data (**Fig. 4a,b, Supplementary Fig. 5a, Supplementary Table 2b**). Microglial programs (*m7_MG1* and *m77_MG2*) were present diffusely throughout the normal brain (**Fig. 4h**), had an early response to sites of injury (injection tract, **Fig 4h**) and persisted long-term within lower density tumor areas (D2-D3) (**Fig. 4f, Supplementary Fig. 5d, Supplementary Table 5e**). MG programs showed differential spatial abundance and distinct pathway enrichments, with *m7_MG1* more abundant overall (**Fig. 4f**), and scoring strongly for IL-1, IL-1b, IL-2 production, for cell migration and response to injury, astrocyte activation, and antigen processing and presentation (**Fig. 4b, Supplementary Table 5a)**. In contrast, *m77_MG2* was enriched for apoptotic cell clearance, and represented a higher percent of all microglia in dense regions (**Fig. 4f**). Two monocyte derived macrophage (MDM) programs (*m40_MDM1*, *m84_MDM2*), were absent from the normal brain but rapidly recruited to early lesions (**Fig. 4h**). These had distinct pathways, with *m40_MDM1* enriched in terms relating to wound and injury response, cell-matrix adhesion, chemotaxis and migration, and *m84_MDM2* scoring highly for macrophage proliferation, positive regulation of phagocytosis, engulfment, and membrane invagination (**Fig. 4b, Supplementary Table 5a)**. Finally, three monocyte programs (*m12_Mo1*, *m55_Mo2*, *m78_Mo3*) were characterized by interferon signaling (Ifit1, Ifit3, Ifit3b, Ifitm3, Ifit2, Ifitm6) (**Fig. 4b, Supplementary Fig.5a, Supplementary Table 5a)**, were spatially co-localized to tumor borders, and did not associate strongly with the early wound response (**Fig. 4h**). They increased in abundance with time, particularly *m78_Mo3* (**Fig. 4h**), which stood out as a proliferating monocyte program also enriched in TNF and IL-1 production**,** thereby linking monocyte inflammatory signaling with the IL-1 response observed in *m59_EndoLV* (**Fig. 4b,d, Supplementary Table 5a).**

Four remaining programs represented immune cell activities (*m20_MDM-ROS*, *m54_ECM-AP*, *m16_Chemokine*, and *m57_Cytotoxic*) (**Fig. 4a,b,i Supplementary Fig. 5a, Supplementary Table 2b**). The most prevalent of these (*m20_MDM-ROS*) had a poor cell type match to MGs/MDMs based on marker genes **(Fig.5i, Supplementary Fig. 5a, Supplementary Table 5b)**, however, spatial localization, pathway activities and TF profiles (**Fig. 4b, Supplementary Fig. 5c, Supplementary Table 5a-d**), indicated this was an MDM activity primarily distinguished by a cellular response to oxidative stress and hypoxia, and regulation of apoptosis and autophagic cell death. *m16_Chemokine* was characterized by immunosuppressive Ccl3 and Ccl4 chemokines[7] (involved in suppression of T/NK cells and in recruitment of Tregs) **(Supplementary Fig. 5a)**. *m54_ECM-AP* was Lyz2 and CD74 high (denoting macrophage activation) and enriched in terms for ECM organization, antigen processing, and transmembrane transport **(Fig. 4b, Supplementary Fig. 5a, Supplementary Table 5a)**. Interestingly, *m16_Chemokine* and *m54_ECM-AP* both co-occurred with MG/MDM programs, but not with each-other, suggesting they are mutually exclusive immune cell activities (**Fig. 5i, Supplementary Fig. 5c**). Finally, *m57_Cytotoxic* was significantly enriched in pathways relating to cell killing, cytokine production and defense response, and found at all timepoints and across tumor densities **(Fig. 4b,f,i, Supplementary Table 5a)**. Relative to other immune cell activities, *m57_Cytotoxic* had the highest prevalence in D4, indicating that cytotoxicity plays a central role in dense regions (**Fig. 4f,i, Supplementary Fig. 5d, Supplementary Table 5e**).

### REFERENCES

1. Heddleston JM, Li Z, McLendon RE, Hjelmeland AB, Rich JN. The hypoxic microenvironment maintains glioblastoma stem cells and promotes reprogramming towards a cancer stem cell phenotype. Cell Cycle [Internet]. 2009 Oct 15 [cited 2023 Nov 26];8(20):3274–84. Available from: https://pubmed.ncbi.nlm.nih.gov/19770585/
2. Li Z, Bao S, Wu Q, Wang H, Eyler C, Sathornsumetee S, et al. Hypoxia-inducible factors regulate tumorigenic capacity of glioma stem cells. Cancer Cell [Internet]. 2009 Jun 2 [cited 2023 Nov 26];15(6):501–13. Available from: https://pubmed.ncbi.nlm.nih.gov/19477429/
3. Puchalski RB, Shah N, Miller J, Dalley R, Nomura SR, Yoon JG, et al. An anatomic transcriptional atlas of human glioblastoma. Science (1979). 2018;360(6389).
4. Das S, Li Z, Noori A, Hyman BT, Serrano-Pozo A. Meta-analysis of mouse transcriptomic studies supports a context-dependent astrocyte reaction in acute CNS injury versus neurodegeneration. J Neuroinflammation. 2020;17(1).
5. Hu X, Deng Q, Ma L, Li Q, Chen Y, Liao Y, et al. Meningeal lymphatic vessels regulate brain tumor drainage and immunity. Cell Research 2020 30:3 [Internet]. 2020 Feb 24 [cited 2023 Nov 26];30(3):229–43. Available from: https://www.nature.com/articles/s41422-020-0287-8
6. Gragnano F, Sperlongano S, Golia E, Natale F, Bianchi R, Crisci M, et al. The Role of von Willebrand Factor in Vascular Inflammation: From Pathogenesis to Targeted Therapy. Mediators Inflamm

[Internet]. 2017 [cited 2023 Nov 26];2017. Available from: https://pubmed.ncbi.nlm.nih.gov/28634421/

7. Ozga AJ, Chow MT, Luster AD. Chemokines and the immune response to cancer. Immunity [Internet]. 2021 [cited 2023 Nov 26];54:859–74. Available from: https://doi.org/10.1016/j.immuni.2021.01.012
